# Integrating host plants and a key natural enemy into MaxEnt improves global suitability predictions for *Semanotus bifasciatus*

**DOI:** 10.1101/2025.10.20.683598

**Authors:** Meiyi Yang, Ya Zou, Yuting Zhou, Yuhang Fan, Shixiang Zong

## Abstract

**BACKGROUND:** *Semanotus bifasciatus* is a major conifer pest that causes severe wood damage. The parasitoid *Sclerodermus guani* is its important natural enemy. However few studies have jointly considered host plants and enemy effects when predicting pest ranges. We applied the optimized MaxEnt model and, on this basis, constructed two models: the “host–pest” model and the “host–pest–enemy” model, to predict the potential global distribution of *S. bifasciatus* under future scenarios and to explore the effects of climate change and the introduction of biotic interactions.

**RESULTS:** Results showed that the climate-only model projected 2.73 × 10□ km² of suitable area under the historical climate condition, concentrated in Asia, North America and Europe, with expansion toward higher latitudes. The expansion of host plants further enhanced pest habitat suitability, nearly doubling the predicted range (5.46 × 10□ km²) and increasing the mean suitability. Moreover, the potential distribution of *S. guani* overlapped extensively with *S. bifasciatus*, reducing the total suitable area of *S. bifasciatus* by up to 2.67 × 10□ km², and mean suitability declined by nearly 40%, indicating effective suppression of pest risk. Centroid shifts were consistently northward, though magnitude and fragmentation varied among models.

**CONCLUSION:** Integrating host availability and enemy suppression improves the realism of distribution forecasts for *S. bifasciatus*. The study highlights the roles of biotic factors in shaping pest suitability and identifies potential future high-risk regions of infestation. These insights provide a solid scientific foundation for targeted monitoring, and the strategic application of biological control in adaptive forest pest management.

## 1 Introduction

*Semanotus bifasciatus* (Coleoptera: Cerambycidae), native to East Asia, is widely distributed in China, Japan, and Korea. It is well known as a destructive wood-boring pest, primarily infesting conifer species such as *Platycladus orientalis*, *Juniperus chinensis*, and *Juniperus rigida* ^1,2^. Its host plants are not only widely used in urban greening and ecological restoration ^3,4^, but also possess significant economic and medicinal value as sources of high-quality timber, industrial raw materials, and internationally recognized herbal remedies ^5,6^. After overwintering inside the host tree, adults emerge in spring, mate, and oviposit in bark crevices, concealing the eggs with secretions ^7^. Neonate larvae bore into the xylem, forming galleries and pupal chambers, and overwinter as adults before emerging from the host. Early infestations cause cortical necrosis, leaf yellowing, and twig dieback; in severe cases, entire trees may die, leading to stand-level mortality ^1,7,8^. Coniferous forests in China, Japan, and Korea have all suffered from *S. bifasciatus*, with China experiencing the most severe and rapidly spreading damage ^2,9^. First recorded in Nanyang, Henan Province in 1992, it had infested nearly one hundred Cupressaceae host trees, causing the death of many individuals by 1999 ^8^. In 1996, due to its devastating impact on ornamental and ancient *Platycladus*, *S. bifasciatus* was designated a forest quarantine pest in China. In 2013, it was officially listed as a nationally significant forestry pest, demonstrating its severity and the urgent need for management ^10^.

Human activities have intensified, leading to an exacerbation of global warming ^11^. According to the Sixth Assessment Report of the Intergovernmental Panel on Climate Change (IPCC), the global temperature increased by 1.09°C (range: 0.95-1.20°C) during the period from 2011 to 2020 compared to 1850–1900. Furthermore, the global average temperature is expected to continue rising over the next two decades, reaching or exceeding 1.5°C above the 1850-1900 levels ^12^. Global warming has clear implications for both fauna and flora ^13^. Insects, being typical ectothermic animals, are directly influenced by temperature and precipitation, which regulate their life history and geographic distribution ^14,15^. Extensive studies have shown that climate change will force insects to move away from unsuitable temperature areas and expand into new regions. It also indirectly affects the multi-trophic interactions between plants, herbivorous insects, and natural enemies. Insect food quality is influenced by changes in plant nutrition and defense compounds, while insect distribution is constrained by the dispersal and adaptability of their host plants ^15,16^. For instance, in Mount Kinabalu, Malaysia, high-altitude moths migrate to higher elevations along with their low-temperature-dependent alpine shrub hosts; in the European Alps and the Pyrenees, stoneflies (Order: Plecoptera) have contracted their distribution to higher altitudes due to their cold-water-dependent hosts (algae and aquatic plants), with 63% of individuals, particularly high-altitude endemics, experiencing range contractions, and some species facing regional extinctions ^15^. The development cycles of natural enemies may shorten in response to climate warming, thereby weakening their effectiveness in pest control ^17^. For example, Barton observed that differing responses to climate warming resulted in reduced spatial overlap between the nursery spider (*Pisaurina mira*) and its prey, the grasshopper (*Melanoplus femurrubrum*), thereby increasing the grasshopper’s feeding opportunities ^18^. Therefore, when predicting the potential geographic distribution of insects under climate change scenarios, it is essential to consider their roles in food webs and the ecological niches they occupy. Incorporating biotic interactions into these predictions makes the results more realistic, and early risk prevention and long-term management strategies based on such predictions will be more valuable.

Currently, over ten species distribution models (SDMs) have been developed for pest risk management and early warning, including the Generalized Linear Model (GLM), Generalized Additive Model (GAM), Random Forest (RF), and MaxEnt. By statistically linking the geographic distribution of species or communities to their current environment, SDMs are commonly used to explain niche shifts and geographic migrations of species ^19,20^. Among the numerous SDMs, the MaxEnt model has become one of the most widely used models for predicting single-species distribution due to its powerful and flexible predictive capabilities and its tolerance for sparse sample data. The MaxEnt model predicts species distributions by calculating the probability distribution of maximum entropy and is considered a purely data-driven model (Elith* et al., 2006; Li et al., 2025). It has demonstrated good performance in both predicting the distribution of single species and averaging the performance across multiple species and regions. Today, MaxEnt is extensively applied in fields such as predicting the potential distribution of insects under climate change, forecasting invasion risk, and delineating protected areas ^23–25^.

Research on *S. bifasciatus* has primarily focused on its biology, chemical ecology, and control strategies. Its wood-boring and concealed lifestyle renders chemical insecticides largely ineffective, highlighting the importance of biological control. Among natural enemies, the ectoparasitic wasp *Sclerodermus guani*, native to East Asia, plays a critical role. It has been widely deployed against cerambycid pests ^26^ and has shown strong efficacy against *S. bifasciatus* in both laboratory and field trials ^27^. In recent years, efforts have been made in China to predict the potential distribution of *S. bifasciatus*; however, most studies have focused solely on climatic drivers, neglecting biotic factors such as host plants and natural enemies, thereby reducing both ecological interpretability and practical applicability.

Therefore, using MaxEnt, this study integrates host plants and the natural enemy *S. guani* into climate-driven species distribution models (SDMs) for *S. bifasciatus* to map global habitat suitability and to assess the effects of climate change, host availability, and enemy pressure. The specific objectives are to:

1. **Establish a climate-only baseline:** fit optimized MaxEnt models with climate and occurrence data for *S. bifasciatus*, its host plants, and *S. guani* to estimate their global suitability.
2. **Assess host effects:** integrate suitability indices of multiple host plants via a harmonic-mean “host–pest” model to predict the distribution of *S. bifasciatus* conditional on host availability.
3. **Quantify natural-enemy regulation:** derive a suppression factor from *S. guani* suitability and the mortality it causes, and incorporate it into a “host–pest–enemy” suppression model to evaluate its regulatory effect on the pest’s potential habitat.

This work addresses a key gap in understanding *S. bifasciatus*–host–enemy interactions under climate change. It provides evidence to guide control and management in forestry, helping to reduce its ecological and economic impacts.

## 2 Materials and methods

### 2.1 Species occurrence data

Global distribution data for *S. bifasciatus*, its host tree species, and *S. guani* were collected from the literature and multiple sources of databases. We identified 11 tree species parasitized by *S. bifasciatus* from the literature. The known global occurrence records for *S. bifasciatus*, the 11 host species, and *S. guani* were obtained from databases such as the Global Biodiversity Information Facility (https://www.gbif.org/occurrence/search), iNaturalist (https://www.inaturalist.org), the Chinese Virtual Herbarium (CVH, http://www.cvh.org.cn), iPlant (https://www.iplant.cn), as well as from literature retrieved via literature retrieval platforms including Web of Science (https://www.webofscience.com) and China National Knowledge Infrastructure (CNKI, https://www.cnki.net). Distribution data for locations that only recorded sampling site names were gathered using the Baidu coordinate picker (https://lbs.baidu.com/maptool/getpoint) and the Amap coordinate picker (https://lbs.amap.com/tools/picker) to approximate their geographic coordinates, which were subsequently verified for accuracy. To ensure the reliability of the distribution data, we used the R package *CoordinateCleaner* ^28^ to clean the data, removing records such as material samples, specimens in museums, or those located in marine areas or around institutions housing captive animals. Additionally, the R package *spThin* ^29^ was used to sparsity the data at 5 km × 5 km grid intervals, ensuring consistency with the 2.5’resolution of the environmental layers. In the end, 195 distribution points for *S. bifasciatus*, 138-9606 distribution points for the 11 host plants, and 112 distribution points for *S. guani* were obtained. See Figure S1 1 for per-species sample sizes.

### 2.2 Filtering and processing of environmental variables

The environmental variables used in this study were obtained from WorldClim 2.1 (http://www.worldclim.org). We downloaded 19 bioclimatic variables and elevation data at a resolution of 2.5 minutes for the current period (1970-2000), and future periods, including the 2050s (2041-2060), 2070s (2061-2080), and 2090s (2081-2100). Future climate conditions were based on data from the Chinese Climate Center’s climate system model (BCC-CSM2-MR) of the Coupled Model Intercomparison Project Phase 6 (CMIP6), in accordance with the native distribution of *S. bifasciatus* and *S. guani*. In this study, we considered three Shared Socioeconomic Pathways (SSPs) for emission scenarios: SSP1-2.6, with the strictest carbon emission controls; SSP2-4.5, with moderate emissions and lower carbon use; and SSP5-8.5, with very high emissions and carbon use, to compare habitat suitability under different emission scenarios. Since elevation is a key factor influencing plant survival ^30^, it was also included in the host plant distribution model predictions. To avoid multicollinearity between environmental variables that affect model accuracy, we performed Principal Component Analysis (PCA) and Pearson correlation analysis using the R packages *FactoMineR* ^31^ and *psych* ^32^. Environmental factors that met the following criteria were retained: cos2 > 0.85 in the principal components and contributions higher than the average, correlations between environmental variables with | r | < 0.85, and a contribution rate ≥ 0.5 in the initial model results. For each species, 5-7 environmental variables were retained for model optimization and final prediction (Table 1).

**Table 1.**
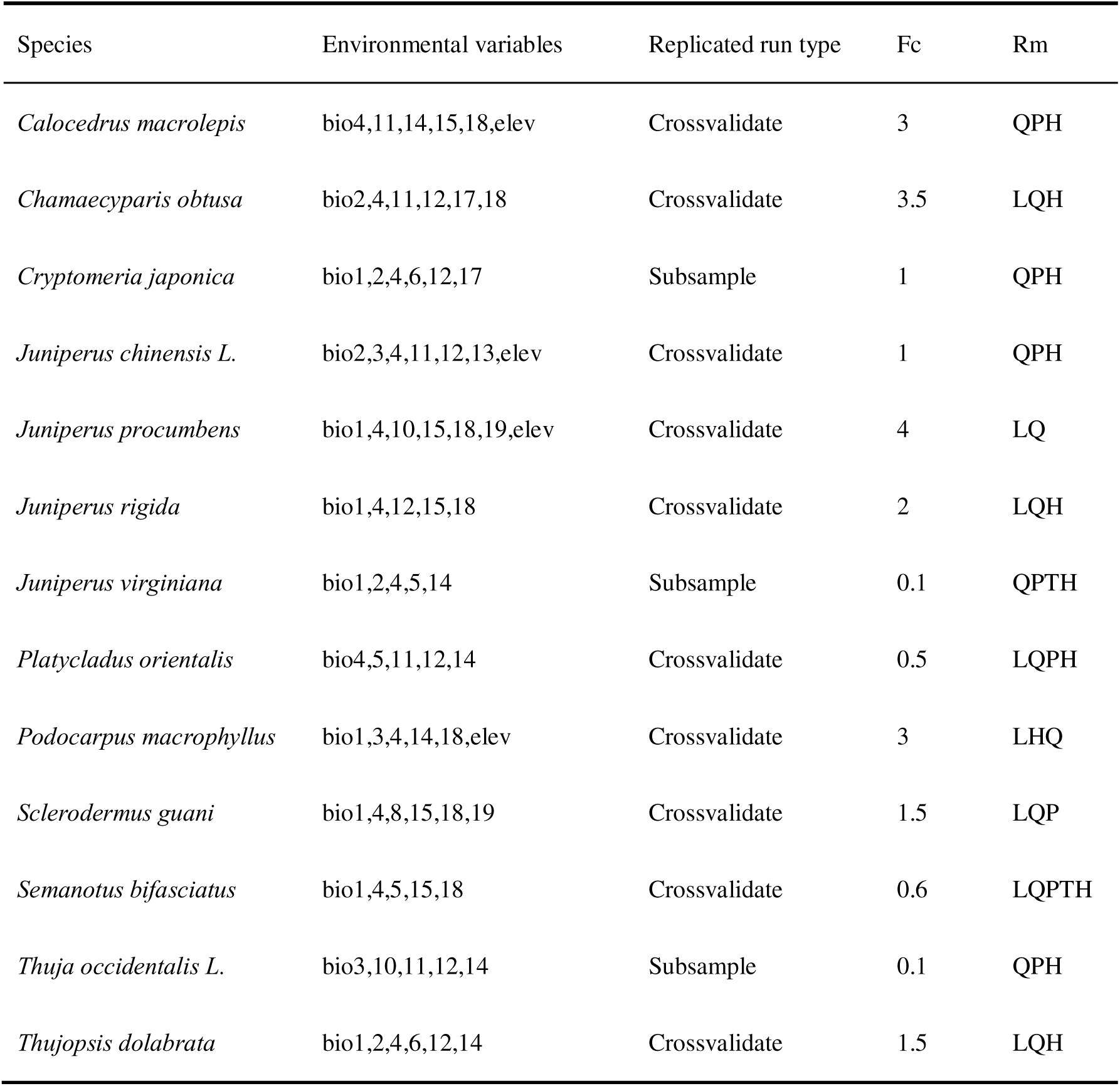
Optimal MaxEnt model settings for each species after tuning, including selected environmental variables, replication type, feature classes (Fc), and regularization multipliers (Rm).

### 2.3 Model optimization and accuracy evaluation of the MaxEnt model

The default settings of MaxEnt can sometimes lead to overfitting of noise in the data and poor adaptability when new conditions are introduced. Therefore, parameter optimization is necessary to enhance the model’s generalization ability. The optimization of the MaxEnt model is achieved through the R package *ENMeval*. *ENMeval* evaluates the model’s performance through cross-validation and selects the optimal parameter combination to enhance prediction accuracy ^33^.

The regularization multiplier (RM) and feature combination (FC) are key parameters in MaxEnt that control the model’s complexity and the use of features. Feature combinations include five main feature types: Linear features (L), Quadratic features (Q), Hinge features (H), Product features (P), and Threshold features (T) ^34^. We set the regularization multiplier within a range of 0.1-4 with an interval of 0.5 and selected eight feature combinations: “L”, “LQ”, “LQP”, “QHP”, “LQH”, “QHPT”, “LQHP”, and “LQHPT”, testing 64 parameter combinations: 8 feature combinations × 8 regularization multipliers. The model with the lowest AICc (Akaike Information Criterion corrected) value (i.e., ΔAICc = 0) was considered the optimal model. When the optimized model results were still not sufficiently superior, the interval for the regularization multiplier was narrowed to 0.2, and the process was repeated until the model met the accuracy evaluation criteria selected for this study.

We used the Test Area Under the Curve (AUC_Test_), Training Omission, and True Skill Statistic (TSS) to evaluate the model’s perefermance. AUC_Test_ demonstrates the model’s performance across different thresholds, ranging from 0 to 1, where a value closer to 1 indicates better model performance. ORMTP (Minimum Training Presence omission rate) and OR10 (10% training omission rate) are threshold-dependent and threshold-related indicators. ORMTP = 0 and OR10 less than 10% generally indicate that the model passes validation ^34,35^. The True Skills Statistic (TSS) measures model performance through a weighted combination of sensitivity and specificity. TSS value ranges from -1 to +1, with 0.4 ≤ TSS ≤ 0.8 indicating an acceptable model, and 0.8 ≤ TSS < 1 considered excellent performance ^36^.

### 2.4 Statistical analysis of the data of the MaxEnt optimal model

#### 2.4.1 Classification of Potential Distribution

The potential distribution prediction results for *S. bifasciatus* were processed using ArcGIS (10.8). For experimental standardization, the natural breaks method was selected to classify the habitat suitability of all models constructed in this study. The global land area was divided into four areas: high-suitability, moderate-suitability, low-suitability, and non-suitable areas. The area calculation was performed using the Raster Calculator in ArcToolbox, and visualization of different areas was carried out in R (4.3.3). For analyzing changes in suitable areas under current and future conditions, we used the “suitability status change index” (SSCI) ^37^. After binarizing the suitable and non-suitable areas based on current and future climate conditions, the areas were categorized into four types: **“**no longer suitable area” (currently suitable areas but that will not remain so in the future), “remaining an unsuitable area” (unsuitable currently and under future climatic conditions), “remaining a suitable area” (suitable currently and under future conditions), and “new suitable areas” (currently unsuitable but becoming suitable under future conditions). The binary threshold was set according to the cutoff between low-suitability and non-suitable areas.

#### 2.4.2 Changes in latitudinal range and centroid migration of potential distribution

The changes in the northern and southern boundaries of the suitable areas, as well as the migration of centroids, were used to quantify the trend of changes in suitable areas. First, the continents with predicted distributions were clipped from the binarized raster data The southernmost and northernmost grid cells with suitability indices ≥ the threshold were extracted for each continent.

Then, the geographical centroid coordinates for each continent under historical climate conditions and future scenarios of different SSPs were calculated, and migration trajectories were plotted. The displacement distance and direction of the centroids relative to current climate conditions were computed. The centroid migration analysis of the suitable areas was completed in ArcGIS (10.8).

#### 2.4.3 Incorporating Biotic Interactions into Predictions

In R, the maximum value of the prediction results for each of the 11 host plant species was extracted for each grid cell to construct the host plant complex. To account for the host factor in the model, we referred to the method used by Zou et al. ^38^..We employed the harmonic mean to construct a distribution prediction model for *S. bifasciatus* that incorporates host plants, as shown in Equation (1). The core advantage of the harmonic mean lies in quantifying the “joint limitation” effect of multiple factors, reflecting the strong dependence of the pest on both host plants and climate. This model satisfies the condition where the insect cannot stabilize its population if only the climate is suitable but no hosts are present (b ≈ 0), or if hosts are abundant but the climate is unsuitable (a ≈ 0). Equation (1) is as follows:

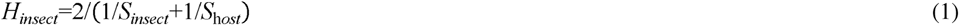

where *H_insect_* represents the harmonic mean suitability index for *S. bifasciatus*, *S_insect_* represents the suitability index for *S. bifasciatus*, and *S_host_* represents the suitability index for the host plants complex.

Building on this, the natural enemy effect was further introduced by setting up a suppression factor, and a dual-impact SDM model was constructed, incorporating both host plants and *S. guani*. In nature, the parasitism rate and mortality rate of the parasitoid are influenced by climate change ^24,39^. This is reflected in the underlying logic of constructing the suppression factor.

In the first step, parasitism data from several previous studies on *S. guani* against *S. bifasciatus* were synthesized to quantify the pest control effect of the natural enemy. According to the literature, under suitable conditions, indoor experiments have shown a mortality rate of over 80% for *S. bifasciatus* larvae after 90 days, while outdoor experiments have shown a mortality rate of approximately 70% ^40–42^. This study took 70% as the control efficacy of *S. guani* under outdoor artificial release conditions. A suppression factor was developed by integrating the suitability index of *S. guani* with empirically derived field parasitism efficacy, defined as Equation (2):

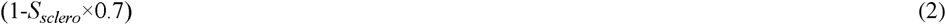

where *S_sclero_* represents the suitability index for *S. guani*.

Finally, the suppression factor was integrated into the previously constructed harmonic mean model to obtain the complete natural enemy suppression model, as shown in Equation (3):

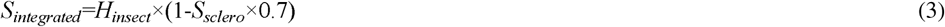

where *S_integrated_* represents the suitability index of the natural enemy suppression model.

#### 2.4.4 Comparison of Model Results

The output results of the three models were subjected to multidimensional quantitative comparison and analysis in R to reveal how biotic factors (host plants, *S. guani*) affect the potential suitable areas of *S. bifasciatus* and to examine the dynamic differences in their impact strength under different time and climate change scenarios. Specifically, we conducted analyses of suitability mean (meanSuit), suitability standard deviation (sdSuit), as shown in Equations (4) and (5), and suitability overlap (Schoener’s D), as shown in Equations (6) and (7).

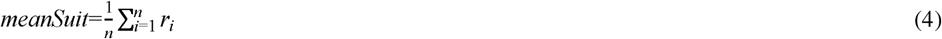

where *r_i_* represents the suitability value in the *i*-th grid cell, and *n* is the number of valid cells, excluding missing data.

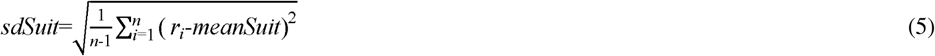

where *r_i_* and *n* are defined as above.

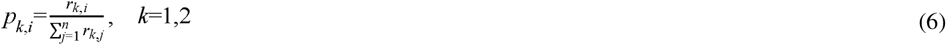

where *r_k,i_* is the suitability value of species *k* at grid cell *i*, *n* is the total number of grid cells, and *p_k,i_*, *i* represents the normalized probability for species *k*.

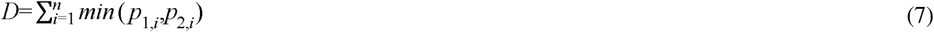

where *p_1,i_* and *p_2,i_* are the normalized probabilities of the two species at grid cell *i*, and *D* ranges from 0 (no overlap) to 1 (complete overlap).

The suitability mean (meanSuit) analysis quantifies the differences in the overall suitability intensity of *S. bifasciatus* across the three models under different time and climate scenarios by calculating the mean suitability of each model. By comparing the mean differences in the results and the percentage of increase or decrease, we assessed the promotion or inhibition of suitability intensity by host plants and *S. guani*. The suitability standard deviation (sdSuit) analysis calculates the standard deviation of suitability in each model to characterize the spatial heterogeneity of suitability within suitable areas. By comparing the differences in standard deviations between models, we revealed the impact of host plants and *S. guani* on the spatial heterogeneity of the suitable areas. We also calculated the suitability overlap (Schoener’s D) between different models based on the spatial matching of raster data to evaluate the spatial congruence between the suitable areas of *S. bifasciatus* and its host plants, and to verify whether the suitable areas of *S. bifasciatus* and its natural enemies are spatially associated.

## 3 Results

### 3.1 Model optimization and accuracy evaluation

Climate variable selection and parameter tuning were performed for the MaxEnt models of *S. bifasciatus*, 11 host plant species, and *S. guani*. The performance of the optimized models improved significantly. Taking *S. bifasciatus* and *S. guani* as examples, their optimal models were obtained when the FC and RM were set to QPH and 3, LQP and 4, respectively. The ΔAICc values for both models were 0, with AUC_test_ values of 0.987 and 0.984, indicating very high predictive accuracy. The omission rate indicators also performed well, with ORMTP = 0 and OR10 values of 0.0977 and 0.0991, respectively. The TSS values of both optimal models were greater than 0.9, confirming their high reliability. The optimized MaxEnt model of *S. bifasciatus*, using the screened variables, showed high predictive performance, laying a solid foundation for the development of reliable new distribution models incorporating biotic factors (Fig. 1, Fig. 2). Relative contributions of environmental variables to the optimized MaxEnt models and their response types for each species is provided in Table S3. Each species summary of the selected evaluation metrics (AUC_Test_, ORMTP, OR10) is provided in Table S3. The contribution and importance of bioclimatic variables for each species are provided in Figure S1 and Figure S2.

**Fig. 1.**
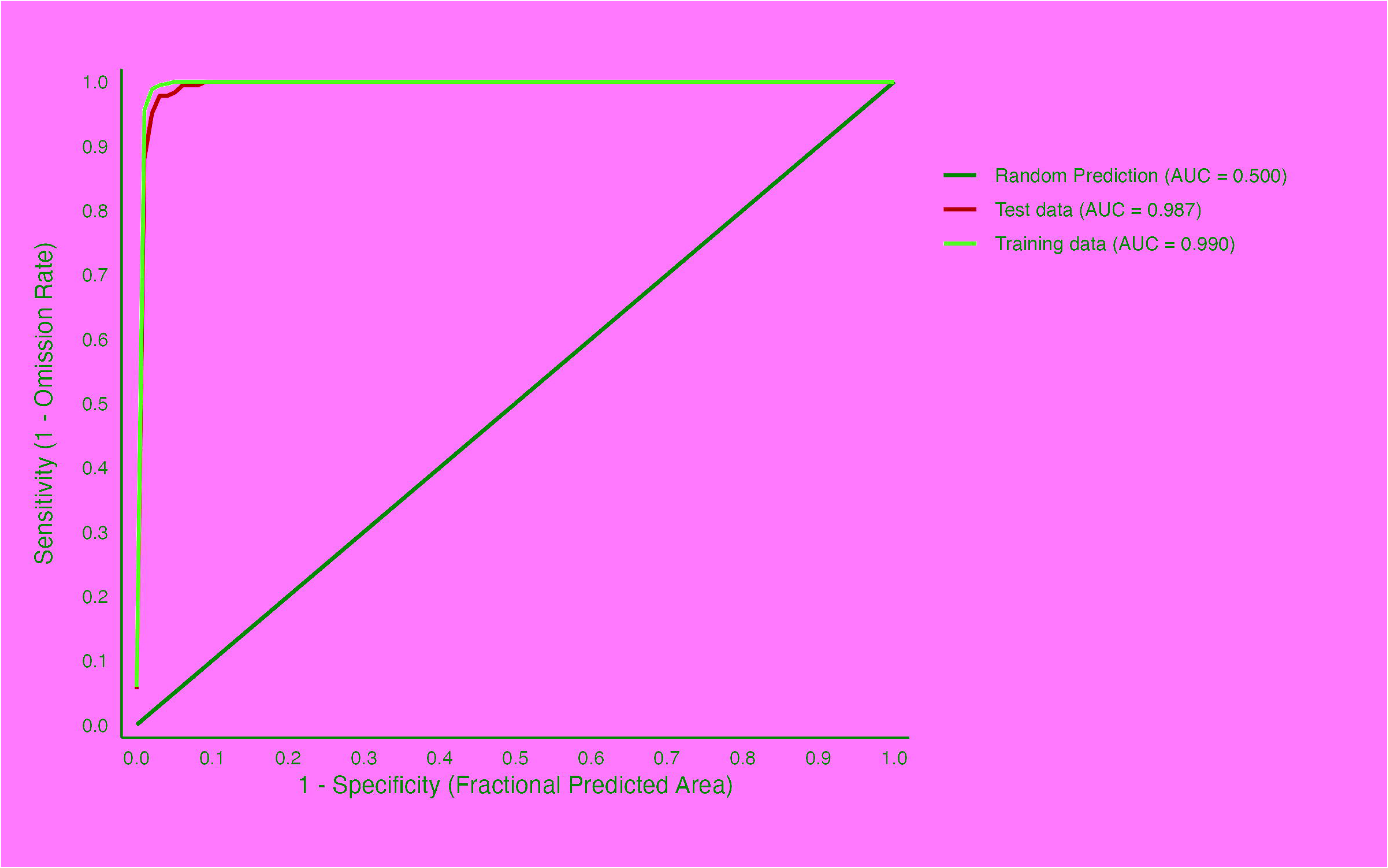
Receiver operating characteristic (ROC) curves of the optimized MaxEnt model for *Semanotus bifasciatus* based on 10 replicates The plot displays receiver operating characteristic (ROC) curves for random prediction (black line, AUC = 0.500), model performance on test data (blue line, AUC = 0.987), and model performance on training data (red line, AUC = 0.990). The x-axis represents 1−Specificity (Fractional Predicted Area), and the y-axis represents Sensitivity (1 - Omission Rate).

**Fig. 2.**
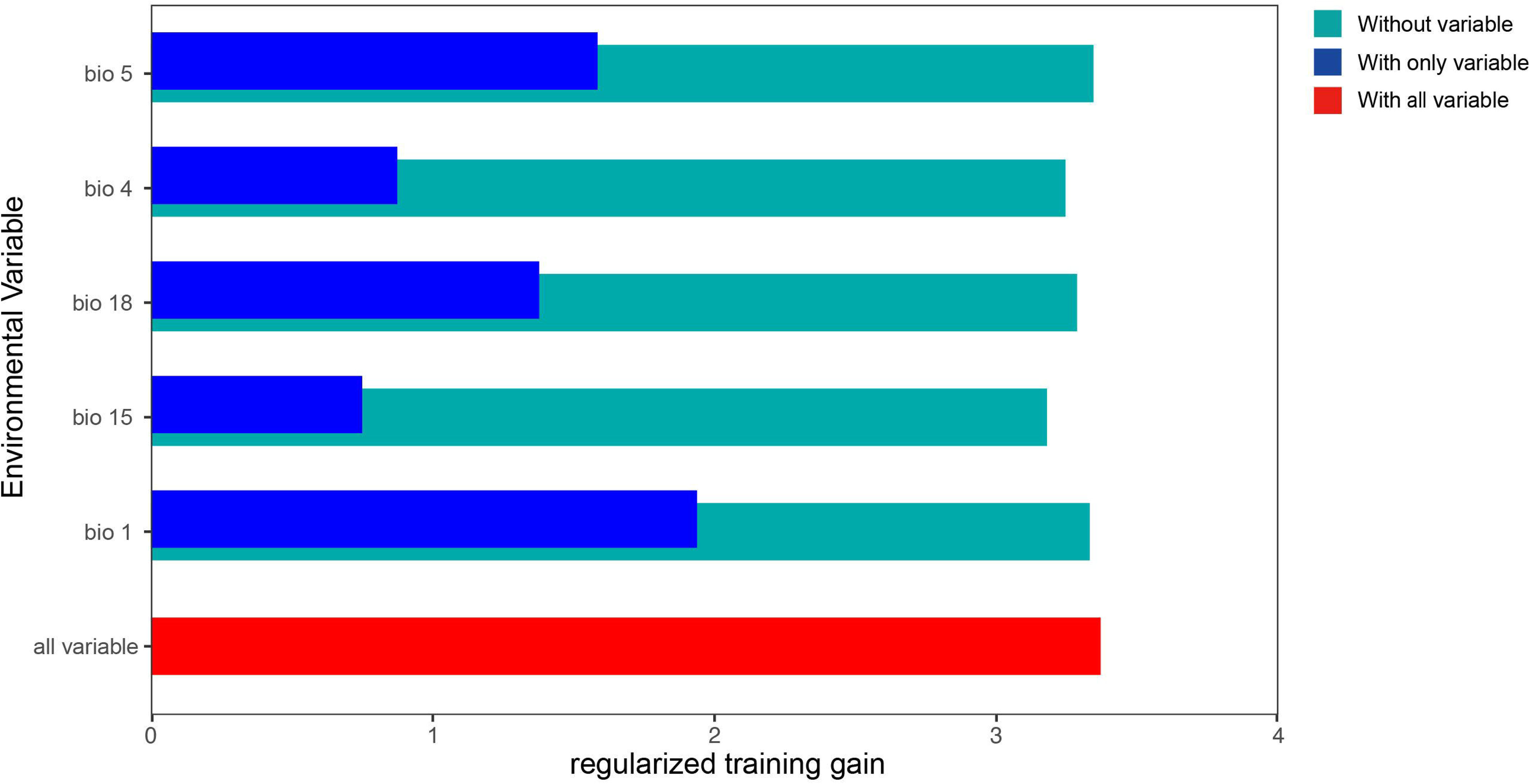
Jackknife test of variable importance for the optimized MaxEnt model of *Semanotus bifasciatus* Teal bars indicate the regularized training gain when the corresponding single variable is excluded, blue bars indicate the gain when only the corresponding single variable is used, red bar indicates the gain when all variables are used.

### 3.2 Prediction of the Potential Distribution of *S. bifasciatus*

#### 3.2.1 Potential distribution under historical climatic conditions

Under current climatic conditions, the total global suitable area of *S. bifasciatus* is approximately 2.73 × 10^6^ km², accounting for about 1.87% of the total global land area and is distributed across three major continents in the Northern Hemisphere: Asia, North America, and Europe. The low-suitability area is the largest, totaling 1.51 × 10^6^ km², which represents 55.12% of the total suitable area. Meanwhile, the moderate- and high-suitability areas account for 24.92% and 19.96%, respectively. All high-suitability areas and the vast majority of moderate-suitability areas are concentrated in eastern Asia, covering China, North Korea, South Korea, and Japan, with China encompassing all three suitability levels. In North America, patchy low-suitability areas are distributed across the central Great Plains of the United States, covering an area of approximately 0.47 × 10^5^ km². The current potential distribution of *S. bifasciatus*, as predicted by the three models, is illustrated in Fig. 3, showing spatial patterns of low-suitability moderate-suitability, and high-suitability areas under historical climatic conditions. Figure S3-S5 illustrate the potential distribution of *S. bifasciatus,* its 11 host plants and *S. guani* under historical and future climates, Figure S6-S8 illustrate the potential suitable area of *S. bifasciatus* under historical and future climates predicted by three models.

**Fig. 3.**
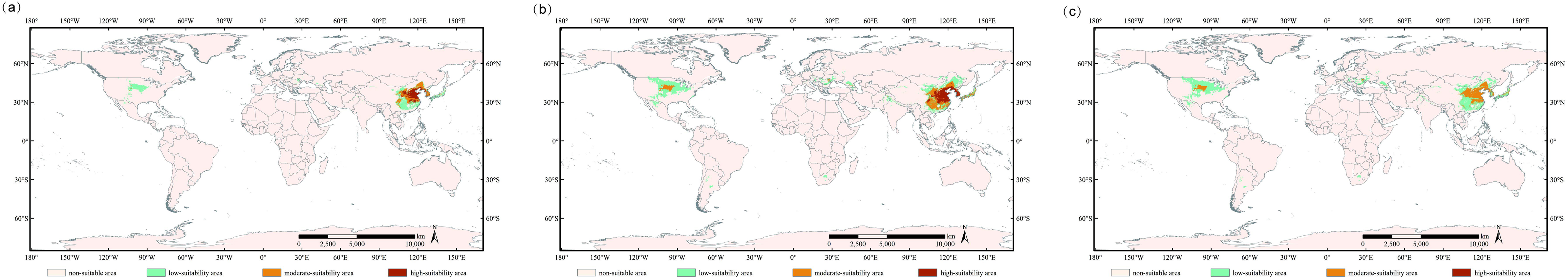
Current potential distribution of *Semanotus bifasciatus* predicted by the three models climate-only model (a), harmonic mean model (b), and suppression model (c) Beige indicates “non-suitable area”, green indicates “low-suitability area”, orange indicates “moderate-suitability area”, and red indicates “high-suitability area”.

#### 3.2.2 Potential distribution under future climatic conditions

Under different SSPs, the total potential suitable area of *S. bifasciatus* generally exhibited an expansion trend, but with markedly distinct migration trajectories. the trend from the 2050s to the 2070s and then to the 2090s followed a “first expansion – subsequent contraction – followed by recovery” curve. In contrast, under SSP5-8.5, the potential suitable area expanded dramatically, increasing by 258.6% relative to the current climate by the 2090s and reaching 9.80 × 10^6^ km². Moreover, the suitability-class structure underwent a fundamental transformation: the proportion of low-suitability areas surged to 70%, while the area of high-suitability habitat sharply declined to only 5.40% of the total (Fig 4). Areas of of low-, moderate-, and high-suitability for each model, scenario, and period are provided in Table S4.

**Fig. 4.**
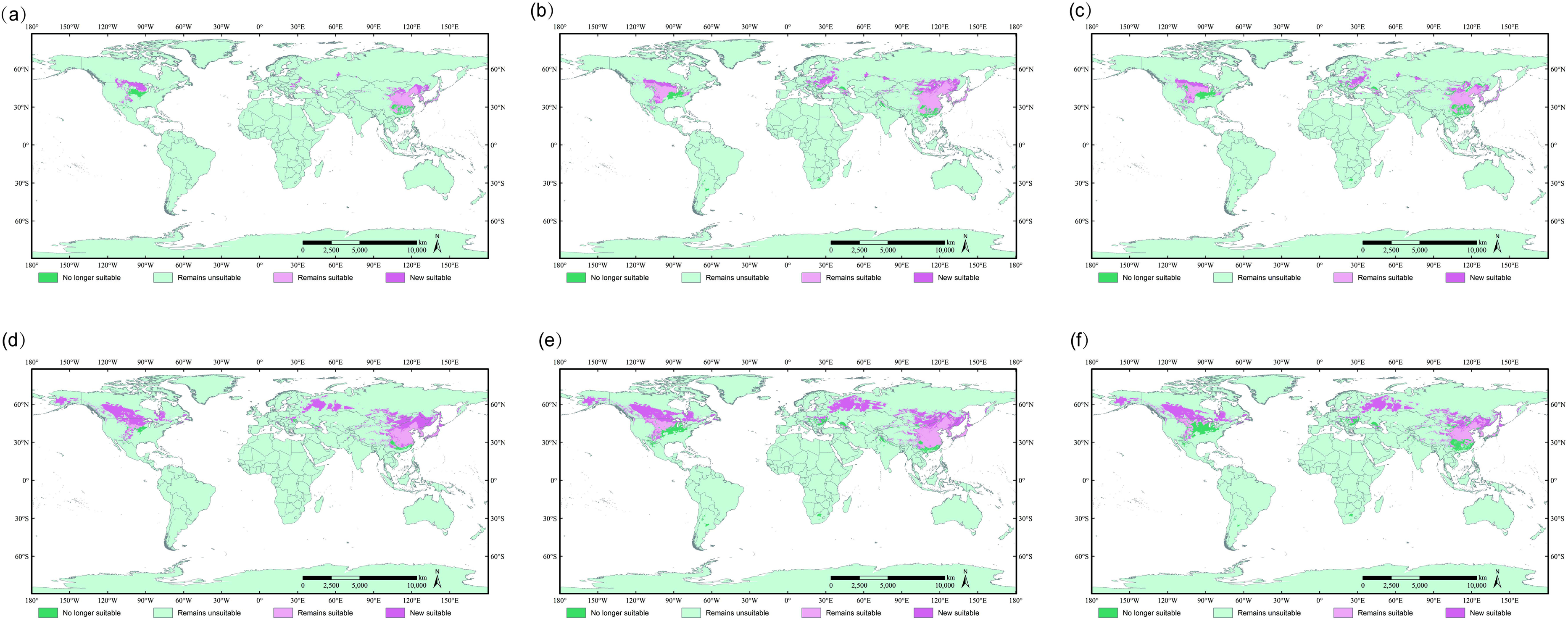
Predicted dynamics of suitable areas for *Semanotus bifasciatus* under SSP1-2.6 (a–c) and SSP5-8.5 (d–f) scenarios by the climate-only MaxEnt model, harmonic mean model, and suppression model Dark green represents “no longer suitable”, light green represents “remains unsuitable”, light pink represents “remains suitable”, and dark pink represents “new suitable”. Panels (a–c) show predictions under SSP1-2.6, while panels (d–f) correspond to SSP5-8.5.

#### 3.2.3 Centroid migration and Spatial pattern changes

Under different scenarios, the overall centroids of suitable areas in Asia and North America shift northwestward. At the same time, in Europe, the centroid migrate westward under SSP1-2.6, and eastward under SSP2-4.5 and SSP5-8.5. Detailed centroid coordinates and migration distances for all models and continents are provided in Table S5. Currently, the centroid of the suitable habitat in Asia is located in Henan Province, China (114.21°E, 35.80°N), and by the 2090s, it is projected to move into Inner Mongolia. Under SSP5-8.5, the high-suitability areas in Asia become severely fragmented (Fig. 5). In North America, the present centroid is situated in New Mexico, United States (-103.87°W, 36.98°N). With a marked increase in suitability, the centroid shifts northwestward under all climate scenarios, reaching up to 16° higher in latitude, and under SSP5-8.5, moderate-suitability areas emerge in southern Alaska and Newfoundland. In Europe, the current centroid lies in Ukraine (35.10°E, 45.27°N). Under SSP1-2.6, the centroid shifts westward into Moldova, with only minor movement of the overall potential distribution. In contrast, under the high-emission scenario, the centroid undergoes a pronounced northward shift into Russia by the end of this century, resulting in the disappearance of the current potential distribution, and new patches of moderate-suitability areas emerge across the Eurasian continent.

**Fig 5.**
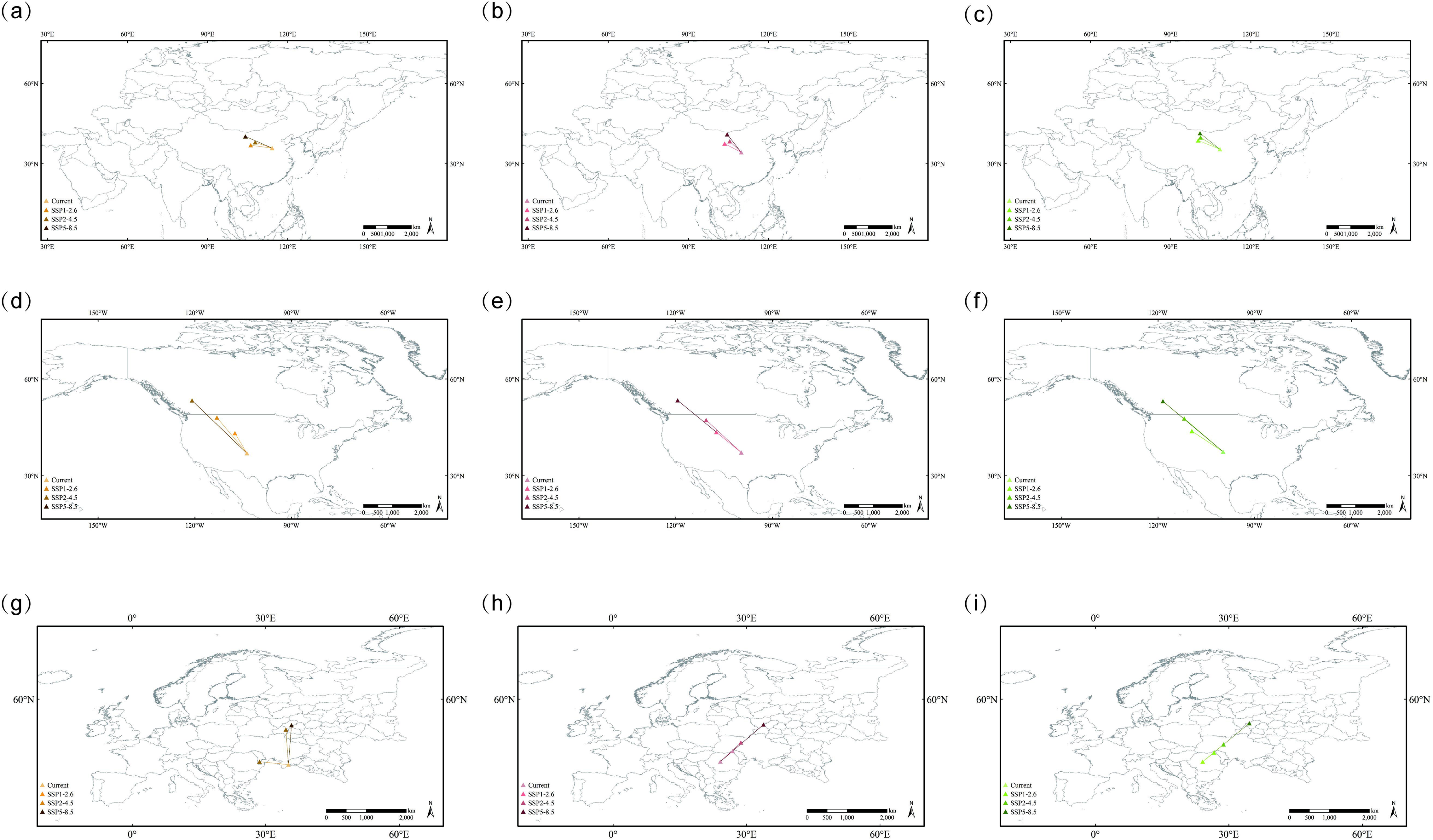
Centroid shifts of suitable areas for *Semanotus bifasciatus* from the current period to the 2090s under SSP1-2.6, SSP2-4.5, and SSP5-8.5 scenarios in Asia (a–c), North America (d–f), and Europe (g–i) Each arrow links the current centroid with its projected location in the 2090s under the three scenarios for the climate-only, harmonic mean, and suppression models.

### 3.3 Prediction of the potential distribution of *S. bifasciatus* considering host plants under climate change

#### 3.3.1 Potential distribution of *S. bifasciatus* under historical climatic conditions

Under historical climate conditions, the global suitable area of *S. bifasciatus* reached 5.46 × 10^6^ km², accounting for approximately 3.7% of the total global land area. The low-suitability area covered about 3.26 × 10^6^ km², constituting the dominant distribution pattern with a proportion of 60%. The moderate- and high-suitability areas were 1.31 × 10^6^ km² and 8.89 × 10^5^ km², respectively, exhibiting a gradient distribution pattern. Asia remained the global core of potential distribution, with a total suitable area of 4.2 × 10^5^ km², accounting for about 9.4% of the continent’s land area.

#### 3.3.2 Potential distribution of *S. bifasciatus* under future climatic conditions

Compared with the predicted results under historical climate conditions, the total global potential suitable habitat expanded the least under SSP1-2.6, increasing by approximately 1.86 × 10^6^km² by the end of this century. Under SSP5-8.5, the expansion was most pronounced, with the total area reaching 1.25 × 10^7^ km² by the end of the century. The increase was mainly in the moderate-suitability area, with the added area exceeding twice the total historical moderate-suitability area, while the high-suitability area shrank to 6.76 × 10^5^ km² by the 2090s.

#### 3.3.3 Centroid migration and Spatial pattern changes

Compared with the climate-only model, the migration trends under different emission scenarios were more consistent. Under SSP5-8.5, extreme warming drove a significant spatial reconfiguration of the global potential area of *S. bifasciatus*. In Asia, the centroid remained within China by the end of the century, shifting northwestward from Shanxi Province to as far as Inner Mongolia. However, the moderate-suitability area migrated markedly to higher latitudes, with high-suitability habitats emerging in Russia and Mongolia. In North America, the expansion of low-suitability areas was pronounced, with centroid migration distances exceeding those in Asia. By the 2090s, the centroid is projected to shift from its current location in Kansas, USA, to Alberta, Canada. In Europe, the centroid consistently migrated northeastward under all scenarios, moving from Romania to ultimately reach Spass-Demensk, Russia.

### 3.4 Prediction of the potential distribution of *S. bifasciatus* considering host plants and *S. guani* under climate change

#### 3.4.1 Potential distribution under historical climatic conditions

Using the same threshold as in the harmonic mean model, the prediction results of the suppression model were classified into suitability levels. The predicted global potential distribution area reached 4.82×10^6^ km², with low-suitability areas accounting for 70.95% and moderate-suitability areas for 29.05%. High-suitability areas in East Asia completely disappeared, being replaced by moderate-suitability areas. Moreover, the moderate-suitability areas in southern China (south of the Yangtze River) and Japan have degraded into low-suitability areas. In contrast, the distribution and extent of moderate- and low-suitability areas in North America and Europe remained nearly unchanged.

#### 3.4.2 Potential distribution under future climatic conditions

*S. guani* exerted a pronounced controlling effect on the spread of *S. bifasciatus*. Under suppression by *S. guani*, the global potential distribution of *S. bifasciatus* across all radiation scenarios and periods consisted solely of low- and moderate-suitability areas, with low- suitability predominating. The minimum suitable distribution occurred under SSP1-2.6 in the 2070s, with a total potential area of 4.99×10^5^ km², nearly identical to the present extent. Assuming continuous and well-regulated biological control in the suitability areas of *S. bifasciatus*, the proportion of moderate-suitability areas in the total suitable area remained below 20% across all scenarios and periods.

#### 3.4.3 Centroid migration and Spatial pattern changes

The inclusion of *S. guani* suppression did not alter the overall direction of centroid migration. In Asia, the current centroid under the suppression model is located in Gansu, China (108°E, 35.4°N) and remains within Chinese territory under all future climatic scenarios. In North America, the centroid is currently located in Kansas, USA (99.81°W, 37.45°N), but under SSP5-8.5, it shifted northward to Alberta, Canada. Notably, by the 2070s, all low-suitability areas within the United States had declined, although by the 2090s, extensive low-suitability areas had reemerged. In Europe, the current centroid lies in Carpathian Ukraine (24.1°E, 45.94°N), with future northeastward shifts proportional to the intensity of radiative forcing.

### 3.5 Impact of Biotic Interactions on the Potential Habitat of *S. bifasciatus*

#### 3.5.1 Effects of Host Plants on the Potential Distribution of *S. bifasciatus*

Host plants not only expanded the distribution of *S. bifasciatus* but also markedly increased the overall suitability and spatial heterogeneity of suitable areas by supplying resources (Fig. 6). With host plants incorporated, the total suitable area of *S. bifasciatus* under historical conditions increased to 5.46 × 10^6^ km². The meanSuit rose from 0.0091 to 0.0123, and this “facilitation effect” was more substantial under more favorable climatic conditions: under SSP1-2.6 in the 2070s, the mean suitability exhibited the most pronounced increase (by 39.34%) (Fig. 7). The sdSuit increased from 0.0615 to 0.07129, because low-suitability areas expanded widely while moderate- and high-suitability areas became locally concentrated.

**Fig. 6.**
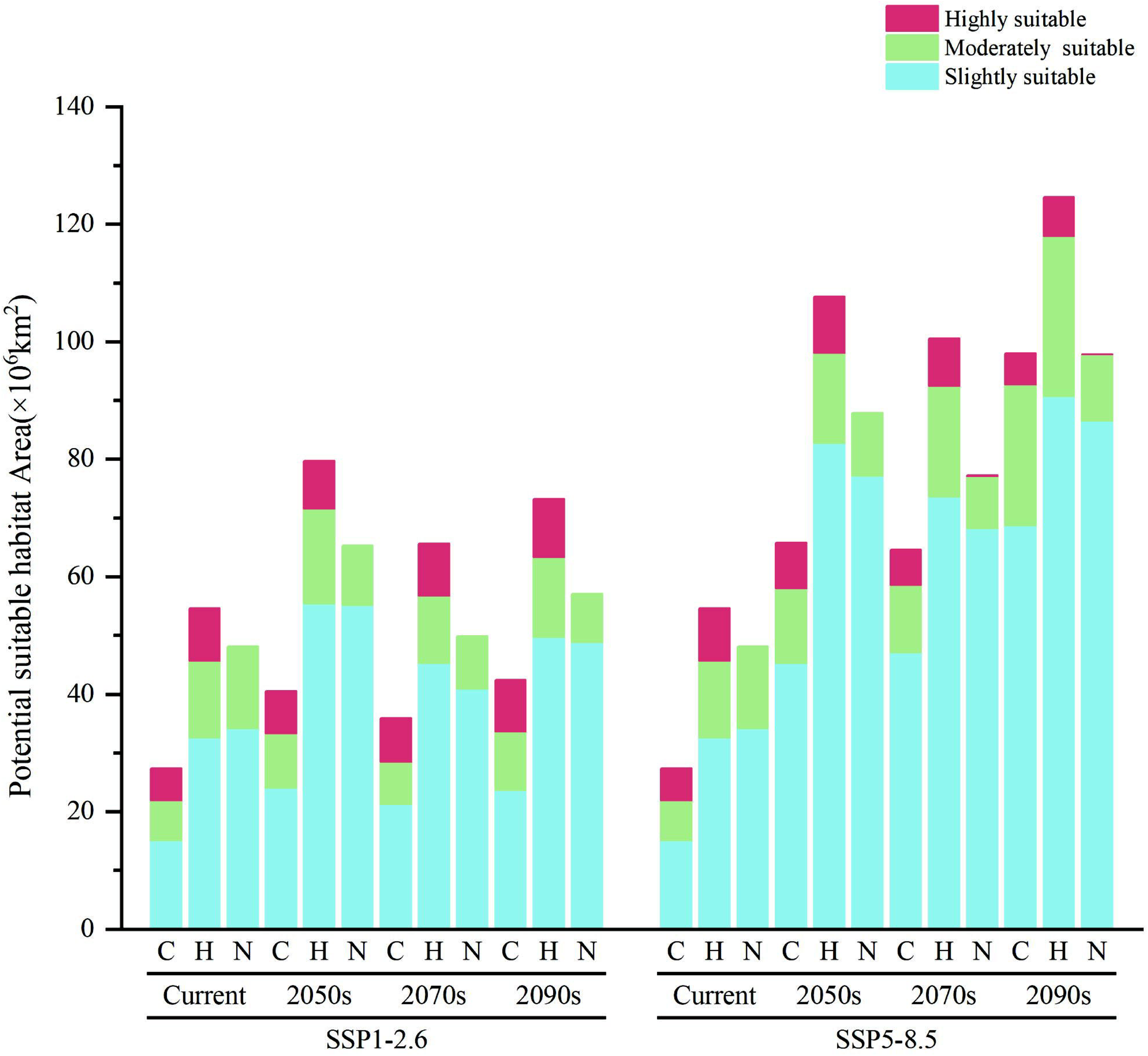
Comparison of potential suitable habitat areas of *Semanotus bifasciatus* among three models across different scenarios and periods Red bars indicate high-suitability areas, green bars indicate moderate-suitability areas, and blue bars indicate low-suitability areas. C, H, and N represent predictions from the climate-only model, harmonic mean model, and suppression model, respectively.

**Fig. 7.**
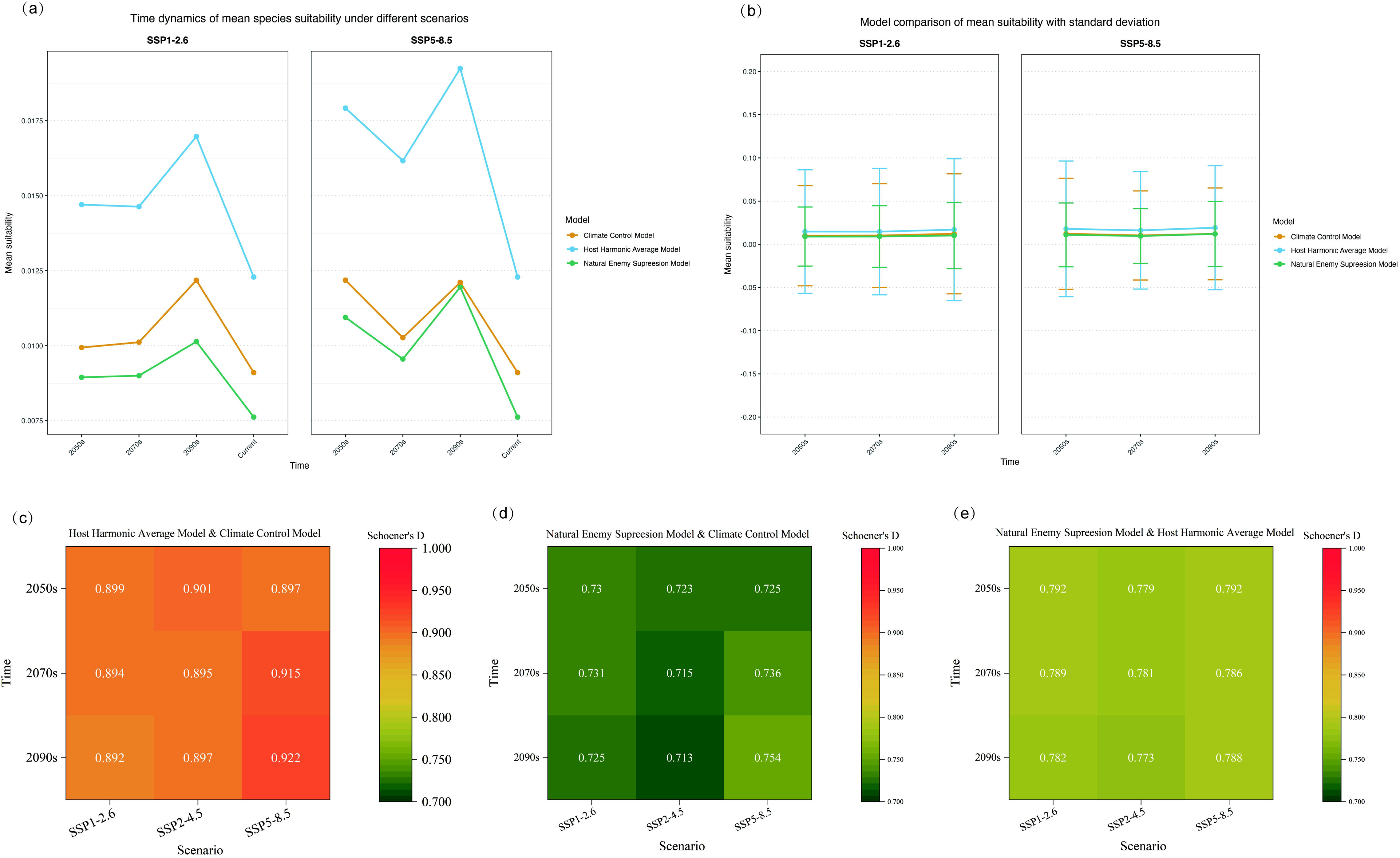
Ensemble comparison of suitability predictions across three models: mean suitability (a), standard deviation (b), and spatial overlap (c) This figure illustrates the concordance and divergence among predictions from the climate-only model, harmonic mean model, and suppression model.

Consistent with this, the potential suitable areas before and after adding host plants showed high spatial congruence, with Schoener’s D remaining high across scenarios (0.8922–0.9216). The greatest overlap occurred under SSP5-8.5 (D = 0.9216), where the difference in total suitable area before vs. after adding hosts was also minimal (only 0.45 × 10^6^ km²). Most of the area gains occurred in low- and moderate-suitability areas, while high-suitability areas increased by only 0.97–2.86 × 10^5^ km².

#### 3.5.2 Regulatory Effects of *S. guani* on the Potential Distribution of *S. bifasciatus*

The control exerted by *S. guani* not only compressed the distribution of *S. bifasciatus* but also rendered the spatial pattern more homogeneous. Under historical conditions, the potential suitable area was 4.82 × 10^6^ km², which was 0.643 × 10^6^ km² lower than the result of the harmonic mean model; the meanSuit declined from 0.0123 to 0.0076, a reduction of 37.99%. Future changes in suitable areas were consistent with the trends in global meanSuit. The sdSuit decreased from 0.0718 to 0.0377, indicating reduced within-area heterogeneity of suitability. This reduction consistent with the increased dominance of low-suitability areas (85–88%).

After accounting for *S. guani,* the suitability overlap with the host-integrated model ranged from 0.7735 to 0.8192 across scenarios, further corroborating the control effect of *S. guani*.

## 4 Discussion

### 4.1 Comparison with Previous SDM and Biotic Interaction Studies

Differences in variable selection and model configuration led to discrepancies between our results and the predictions of suitability and geographic distribution for *S. bifasciatus* reported by Cui et al. (2020). and Sun et al. (2024). The former focused on Shanxi Province, while the latter covered the potential global suitable distribution. Multiple factors account for these differences. First, the variation in original occurrence data and the choices related to data complexity exert a strong influence on model performance ^45^. Second, the setting of suitability probability thresholds also plays a critical role in determining the partitioning of suitability classes ^46^. Cui et al. ^43^ relied on only 20 occurrence records, which resulted in low predicted suitability levels for Shanxi Province. In comparison, our study used 195 records, thereby reducing the risk of underestimating dispersal potential, a common issue when sample sizes are small ^47^. In addition, their study employed the default parameters of the MaxEnt model without screening climate variables, which may have resulted in artificially inflated AUC values. Sun et al.^44^ did not incorporate biotic factors into their model, leading to deviations from the species’ actual ecological requirements; as a result, the correspondence between topography and climate in the predicted potential distribution was inconsistent. By optimizing model parameters and integrating the effects of biotic interactions into the modeling of *S. bifasciatus*, the present study improved the reliability of predictions. It more accurately reflected the potential distribution of the species under future climate change scenarios.

Previous researchers have also explored the roles of host plants and natural enemies using various approaches. For instance, Guo et al. ^48^ combined host occurrence data with climate variables as environmental inputs to predict habitat suitability of *Sphaerolecanium prunastri*. Zhang et al. (2024) employed Schoener’s D to construct an Adaptability Overlap Index (AOI) to evaluate the control effects of two natural enemies of the small white-marmorated longicorn (*Monochamus sutor)*, thereby determining priority regions for the application of natural enemies. However, these methods were limited to single aspect, such as host presence or the suitability of natural enemies, without accounting for how climate change may affect host survival status or the control efficacy of natural enemies (e.g., mortality rates).

### 4.2 Validity of the selected dominant environmental factors

The dominant factors for *S. bifasciatus* were precipitation of the warmest quarter (bio18), annual mean temperature (bio1), temperature seasonality (bio4), and maximum temperature of the warmest month (bio5), which are consistent with previous studies on its biological characteristics and are directly related to the climatic types of the three the three Northern Hemisphere continents (Fig. 2). Although the suitable distribution areas across the three continents belong to different climatic areas, their temperature characteristics and seasonal precipitation patterns are highly similar, characterized by “cold winters, hot summers, and synchronous heat and rainfall”. The wettest quarter coincides with the primary growth season of the species, meeting the requirement of bio18 for its growth and reproduction. Bio1 determines the basic thermal conditions for survival, defining the fundamental temperature range in which the species can persist. Bio4 reflects tolerance to environmental fluctuations, and excessively high bio4 values restrict the species’ distribution. Model predictions indicate that the optimal bio1 range for *S. bifasciatus* is approximately 9–16 °C, with a tolerable limit of around 5–20 °C. The most suitable bio4 range is 985–1154 (standard deviation × 100). On the Mongolian Plateau, winters are long and extremely cold, with projected minimum temperatures of approximately −8 to −11 °C under all radiation scenarios and future periods. In the Siberian Plain, future mean bio4 values remain above 1300. Since the dominant climatic variables in these regions exceed the tolerance thresholds of *S. bifasciatus*, they are spared from invasion risks. Bio5 represents the upper limit of extremely high temperatures. When bio5 exceeds the species’ tolerance threshold, suitable areas become “truncated”. Under SSP5-8.5, severe climate change is projected to increase the maximum monthly temperatures in the native suitable regions in East Asia to over 35 °C, surpassing the species’ tolerable maximum (28.06–31.75 °C). This will result in a substantial decline in habitat suitability and a pronounced northward shift of its overall distribution. Therefore, temperature extremes and climatic variability play a decisive role in defining the survival boundaries of *S. bifasciatus*, serving as the fundamental drivers behind its future range contraction and northward migration.

For *S. guani*, the two dominant environmental factors contributing most to its suitable distribution were precipitation of the warmest quarter (bio18) and temperature seasonality (bio4). Bio4 reflects the species’ tolerance to climatic fluctuations, while bio18 indicates that warm and humid conditions are the two key factors for its survival and reproduction. This was also reflected in the prediction results: the flat terrain of the North American Great Plains facilitates north–south air mass exchange, while the regulation of warm and moist air from the Gulf of Mexico moderates annual temperature fluctuations, resulting in projected bio4 values around 700–1300 in the future. This explains why multiple models consistently predicted that this region could develop into a large area of high habitat suitability. Similarly, the sustained warming and humidifying effects of warm currents create favorable conditions for the east coasts of the three continents in the Southern Hemisphere to become potential suitable areas ^50^. Early studies have shown that the parasitism of *S. guani* is limited by low temperatures, while it demonstrates stronger adaptability to high temperatures ^41,51^. The dominant environmental factors identified in the model are consistent with this finding. Therefore, warm and humid conditions combined with moderate climatic variability constitute the critical determinants for the formation and future expansion of suitable areas for *S. guani*.

Among the 11 host plants, the environmental factors with the highest frequency of occurrence indicated that low winter temperatures and drought stress are the main threats to plant survival. Excessive cold can lead to cell freezing and metabolic disruption, while frozen soils in winter hinder root water uptake, thereby reducing stomatal conductance and nitrogen utilization during photosynthesis ^52^. Therefore, survival constraints caused by low winter temperatures and drought stress are the key factors determining the distribution of host plants and their potential shifts under future climate scenarios.

### 4.3 Species distribution influenced by biological interactions

Although *S. bifasciatus* is primarily distributed in East Asia, its host plants occur across Asia, North America, and Europe. Incorporating host plants into the model significantly expanded the potential suitable area for *S. bifasciatus*. This can be attributed to two main mechanisms. First, host plants regulate microclimatic conditions, and arthropods are capable of tracking these favorable microclimates to establish themselves in optimal areas ^53^. Even when large-scale climatic conditions (e.g., annual mean temperature, maximum temperature of the hottest month) are suboptimal for *S. bifasciatus*, host plants can create relatively stable local microclimates that allow the beetle to complete its life cycle. Studies have shown that in summer, certain positions on tree trunks can be several degrees cooler than the surrounding air ^54^, and internal trunk temperatures fluctuate far less than external air temperatures; the insulating capacity of bark can prevent intense summer heat from penetrating the xylem ^55,56^. Renaud et al. (2011; 2009) reported that mixed forests can reduce daily maximum summer temperatures by approximately 5 °C, whereas coniferous forests can raise winter daily minimum temperatures by about 6 °C. These buffering effects create favorable conditions for adult oviposition and larval development, causing areas previously classified as “unsuitable” due to climate to be reclassified as “low-suitability”, with some low-suitability areas upgraded to “moderate-suitability”. Second, the widespread distribution of host plants provides *S. bifasciatus* with “potential dispersal corridors”. The concept of an “empty niche” in invasion ecology refers to regions that possess the necessary resources and environmental conditions for a species but remain unoccupied due to historical or geographic constraints. The presence of host plants represents resource availability, facilitating the establishment of beetle populations upon dispersal ^59^. Consequently, incorporating host plants enables the model to identify additional potential niches, particularly in regions where climatic conditions approach critical thresholds but hosts are present or high-suitability for them.

Introducing *S. guani* markedly reduced the potential suitable area of *S. bifasciatus*, likely due to the parasitoid’s greater tolerance to environmental conditions. The potential distribution of *S. guani* not only fully overlaps with all potential distributions of *S. bifasciatus* but also extends beyond, particularly in the East Asian monsoon region. Under SSP5-8.5, the expansion of *S. guani’s* suitable area spatially and temporally complements the contraction of *S. bifasciatus’s* suitable area. This provides a robust theoretical foundation for the use of parasitoids in biological control and offers strong empirical support for implementing *S. guani* releases in the potential habitats of *S. bifasciatus*. However, it must be noted that transcontinental introductions of *S. guani* may pose invasion risks; therefore, screening and development of local natural enemy resources are recommended in newly invaded areas.

### 4.4 Implications of Dynamic Changes in Potential Distribution

With future climate warming, the effective accumulated temperature during the insect activity season is expected to increase, and the distributional shift of *S. bifasciatus* will coincide with the upward movement of isotherms, contracting from low-latitude and low-altitude boundaries while expanding toward higher latitudes and altitudes ^13,60^. In the harmonic mean model, projections under SSP1-2.6 indicated the smallest expansion of global suitable areas, suggesting that moderate climate change may effectively curb the risk of species spread. However, under SSP5-8.5, the emergence of moderate-suitability areas at high latitudes implies that *S. bifasciatus* may respond to rapid climate change through adaptive evolution. The strong contraction of high-suitability areas suggests that established dominant populations may migrate or undergo local extinction due to heat stress, with new ecological niches being occupied by either more heat-tolerant genotypes or invasive species.

The geographic endpoints of centroid migration exhibit significant ecological barrier effects, with the overall migration trends of potential suitable areas predicted by different models showing consistency and terminating at similar locations. The spatiotemporal dynamics of the suitable areas of *Platycladus orientalis*, a major host plant in East Asia, align with those of *S. bifasciatus*, posing new threats to forest ecosystems in northern China ^61^. However, the Qinghai–Tibet Plateau acts as a natural dispersal barrier, causing the Asian centroid under SSP5-8.5 to halt in the Alxa Desert region. Similarly, the Rocky Mountains restrict the northwestern expansion of *S. bifasciatus* in North America, where new populations may eventually establish in southern Canada, competing with native insects for ecological niches and threatening local forest resources. In contrast, due to the absence of major mountain barriers, the European centroid is projected to extend deep into the inland East European Plain and low-latitude Siberia, warranting heightened vigilance regarding invasion risks. Taken together, these results demonstrate that geographical barriers, in concert with climate change, critically shape the shifting patterns of species’ suitable habitats.

### 4.5 Future directions

Although *S. bifasciatus* currently remains stable within its native distribution, the expansion and migration trends of its potential suitable areas, coupled with intensified human activities, may pose new ecological and economic crises. First, globalization-driven trade and cross-border transportation of timber and wood products greatly increase the risk of species spread, enabling *S. bifasciatus* to overcome natural geographic barriers and invade new regions. Second, land-use change and urbanization fragment host habitats, potentially leading to an overestimation of habitat suitability. Future studies should integrate factors such as the normalized difference vegetation index (NDVI) and the site quality index to establish more precise and robust predictive models. Incorporating biotic factors into species distribution predictions is undoubtedly indispensable; however, ensuring scientific rigor in the face of complex interspecific interactions remains a challenge. For example, the host preference of *S. bifasciatus* and the resistance levels of different host plants remain unclear. Moreover, host adaptability and beetle suitability may not necessarily show a positive correlation. Thus, the weighting of suitability indices for different host plants, as well as quantitative methods for assessing host adaptability, require further investigation. Reseach showed that have demonstrated that *S. guani* inoculated with white muscardine fungus (*Beauveria bassiana)* can increase the mortality of *S. bifasciatus* by approximately 10%. Additionally, the synergistic control potential of *S. guani* with predatory mites warrants further exploration ^62^. Therefore, future research can be further extended along this direction by constructing multi-stressor models that assess the contribution of “biological missiles” to improved biological control outcomes. Meanwhile, integration of remote sensing, GIS, and other technological approaches will enable real-time monitoring of *S. bifasciatus* distribution and population dynamics, thereby providing a solid scientific basis for precision chemical application and biological control.

## 5 Conclusion

This study, incorporated host plant distribution complexes and a natural enemy suppression into the *S. bifasciatus* model, constructing a “host–pest” harmonic mean model and a “host–pest–enemy” suppression model to analyze the spatiotemporal dynamics of the species’ suitable habitats under global climate change. The results indicate that the distribution of host plants significantly expanded the global potential range of *S. bifasciatus* and accentuated regional differences. At the same time, the introduction of the parasitoid *S. guani* effectively suppressed the pest’s spread and reduced the proportion of high-suitability areas. Under future climate scenarios, the expansion of suitable habitats was smallest under the low-emission scenario. In contrast, the total area surged and the spatial pattern was reshaped under the high-emission scenario. Incorporating host and natural enemy factors yielded predictions more consistent with ecological reality, thereby confirming the potential of biological control in curbing pest spread. This study provides a scientific basis for global risk warning, quarantine policymaking, and the optimization of biological control strategies for *S. bifasciatus*. Furthermore, it deepens the understanding of pest–host–enemy interactions under climate change, offering new perspectives and guidance for the adaptive management of similar xylophagous pests, the delineation of priority control regions, and the optimization of release strategies.

## Acknowledgements

This research did not receive any specific grant from funding agencies in the public, commercial, or not-for-profit sectors. However, the authors acknowledge the support provided by the Beijing Forestry University for supplying the necessary laboratory facilities and computational resources.

