## Supplemental Information for "Integrating host plants and a key natural enemy into MaxEnt improves global suitability predictions for *Semanotus bifasciatus*"

**Table S1:** Global occurrence records of all species used in this study

| Species | Occurrence records (n) | Final sample size used in MaxEnt |
| --- | --- | --- |
| *Calocedrus macrolepis* | 204 | 139 |
| *Chamaecyparis obtusa* | 988 | 582 |
| *Cryptomeria japonica* | 6348 | 1584 |
| *Juniperus chinensis* | 774 | 441 |
| *Juniperus procumbens* | 216 | 145 |
| *Juniperus rigida* | 1522 | 735 |
| *Juniperus virginiana* | 35147 | 9606 |
| *Platycladus orientalis* | 1707 | 1091 |
| *Podocarpus macrophyllus* | 614 | 406 |
| *Sclerodermus guani* | 156 | 112 |
| *S. bifasciatus* | 226 | 195 |
| *Thuja occidentalis* | 24870 | 7065 |
| *Thujopsis dolabrata* | 203 | 138 |

**Table S2 :** Relative contributions of environmental variables to the optimized MaxEnt models and their response types for each species

| Species | Environmental variable | Percent contribution (%) | Permutation importance (%) |
| --- | --- | --- | --- |
| *Calocedrus macrolepis* | bio18 | 69.9 | 68.4 |
|  | bio4 | 11.8 | 5 |
|  | bio15 | 7.7 | 9 |
|  | bio11 | 5.8 | 15.6 |
|  | bio14 | 2.4 | 0.9 |
|  | elevation | 2.4 | 1 |
| *Chamaecyparis obtusa* | bio18 | 55.5 | 7 |
|  | bio11 | 17.7 | 81.8 |
|  | bio17 | 15.8 | 0.2 |
|  | bio4 | 6.1 | 2 |
|  | bio12 | 3.1 | 1.4 |
|  | bio2 | 1.9 | 7.6 |
| *Cryptomeria japonica* | bio17 | 59 | 2.2 |
|  | bio6 | 17.3 | 3.5 |
|  | bio1 | 10 | 79.5 |
|  | bio4 | 6 | 3.2 |
|  | bio2 | 5.5 | 7.2 |
|  | bio12 | 2.1 | 4.5 |
| *Juniperus chinensis* | bio11 | 35.7 | 70.1 |
|  | bio12 | 32.9 | 5.6 |
|  | bio4 | 15.5 | 15 |
|  | bio2 | 9.1 | 8.1 |
|  | elevation | 6.8 | 1.2 |
| *Juniperus procumbens* | bio18 | 39.5 | 11.9 |
|  | bio10 | 26.8 | 30.5 |
|  | bio1 | 23 | 54.2 |
|  | bio5 | 6.6 | 1.4 |
|  | bio4 | 3.9 | 1.9 |
|  | elevation | 0.3 | 0.1 |
| *Juniperus rigida* | bio18 | 67 | 20.7 |
|  | bio4 | 21.9 | 19.4 |
|  | bio1 | 4.3 | 55.9 |
|  | bio15 | 3.8 | 2.5 |
|  | bio9 | 2.4 | 0.4 |
|  | bio12 | 0.4 | 0.8 |
|  | elevation | 0.2 | 0.2 |
| *Juniperus virginiana* | bio14 | 45.5 | 21.6 |
|  | bio1 | 25.5 | 51.1 |
|  | bio4 | 17.1 | 22.7 |
|  | bio5 | 9.5 | 3.2 |
|  | bio2 | 2.4 | 1.5 |
| *Platycladus orientalis* | bio11 | 45.6 | 70.8 |
|  | bio12 | 22.4 | 9.3 |
|  | bio14 | 18 | 4.2 |
|  | bio3 | 8.1 | 5.3 |
|  | bio4 | 5.9 | 10.3 |
| *Podocarpus macrophyllus* | bio18 | 73.0 | 31.3 |
|  | bio4 | 15.3 | 0 |
|  | bio1 | 4.8 | 62.9 |
|  | bio14 | 3.8 | 0.6 |
|  | elevation | 1.9 | 0.3 |
|  | bio3 | 1 | 4.8 |
| *Sclerodermus guani* | bio18 | 52.7 | 24.2 |
|  | bio4 | 20.8 | 18.1 |
|  | bio15 | 11 | 7.1 |
|  | bio8 | 9.3 | 17.6 |
|  | bio1 | 6.2 | 32.9 |
| *Semanotus bifasciatus* | bio18 | 28.1 | 10.6 |
|  | bio1 | 25.9 | 44.6 |
|  | bio4 | 22.2 | 36.6 |
|  | bio15 | 12.4 | 6.9 |
|  | bio5 | 11.4 | 1.3 |
| *Thuja occidentalis* | bio14 | 49.9 | 5.2 |
|  | bio11 | 17.6 | 14.2 |
|  | bio12 | 16.8 | 26.6 |
|  | bio10 | 13.1 | 41.5 |
|  | bio3 | 2.5 | 12.5 |
| *Thujopsis dolabrata* | bio14 | 47 | 1.1 |
|  | bio6 | 26.7 | 8.2 |
|  | bio2 | 10.9 | 18.9 |
|  | bio4 | 5.7 | 0.7 |
|  | bio1 | 5.4 | 67.7 |
|  | bio12 | 4.3 | 3.3 |

**Table S3 :** Summary evaluation of optimized MaxEnt models for all species

| Species | AUC_Test_ | ORMTP | OR10 |
| --- | --- | --- | --- |
| *Calocedrus macrolepis* | 0.9862 | 0 | 0.0959 |
| *Chamaecyparis obtusa* | 0.9688 | 0 | 0.1 |
| *Cryptomeria japonica* | 0.9289 | 0 | 0.0996 |
| *Juniperus chinensis* | 0.9598 | 0 | 0.0998 |
| *Juniperus procumbens* | 0.9726 | 0 | 0.0932 |
| *Juniperus rigida* | 0.9640 | 0 | 0.0989 |
| *Juniperus virginiana* | 0.9316 | 0 | 0.1 |
| *Platycladus orientalis* | 0.9278 | 0 | 0.0998 |
| *Podocarpus macrophyllus* | 0.9709 | 0 | 0.0997 |
| *Sclerodermus guani* | 0.9836 | 0 | 0.0991 |
| *S. bifasciatus* | 0.9866 | 0 | 0.0977 |
| *Thuja occidentalis* | 0.9343 | 0 | 0.0998 |
| *Thujopsis dolabrata* | 0.9774 | 0 | 0.0988 |

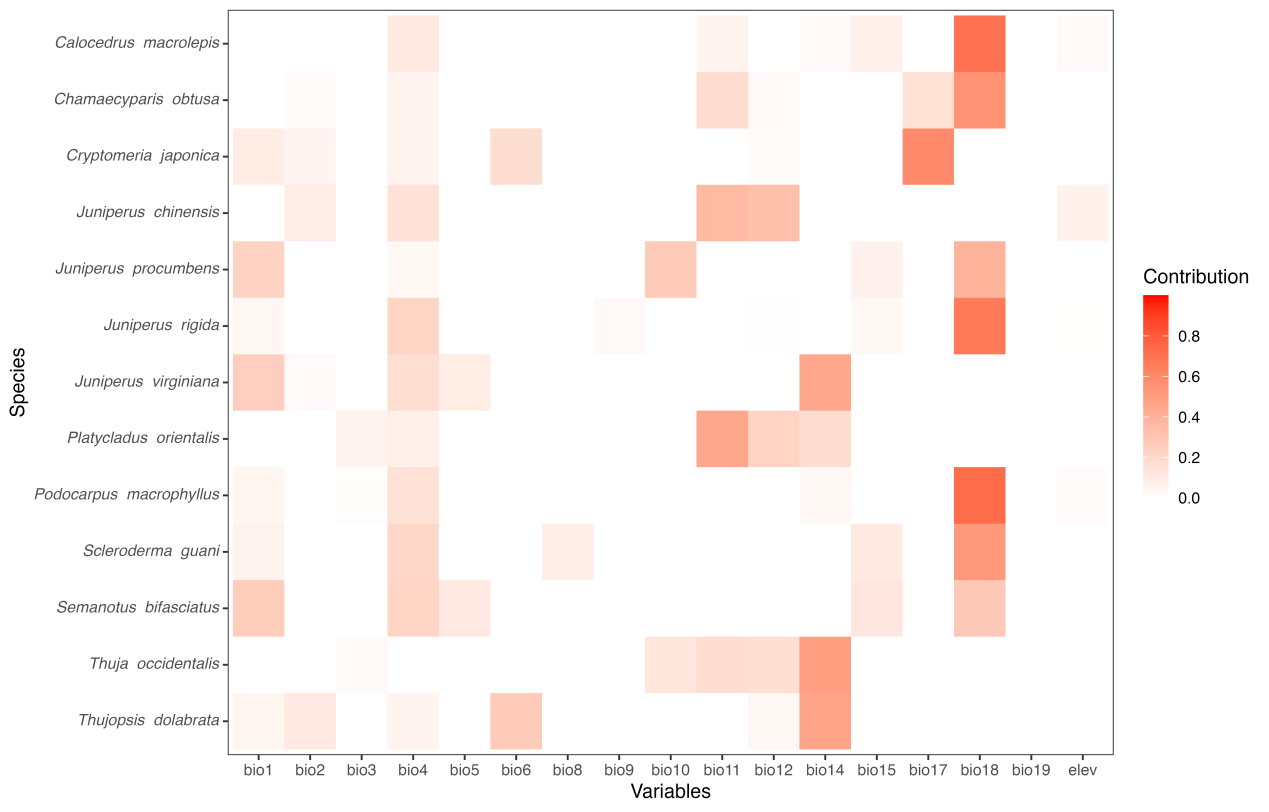

**Figure S1 :** Bioclimatic variable used in the projections for each species.

The red gradient represents the variable contribution.

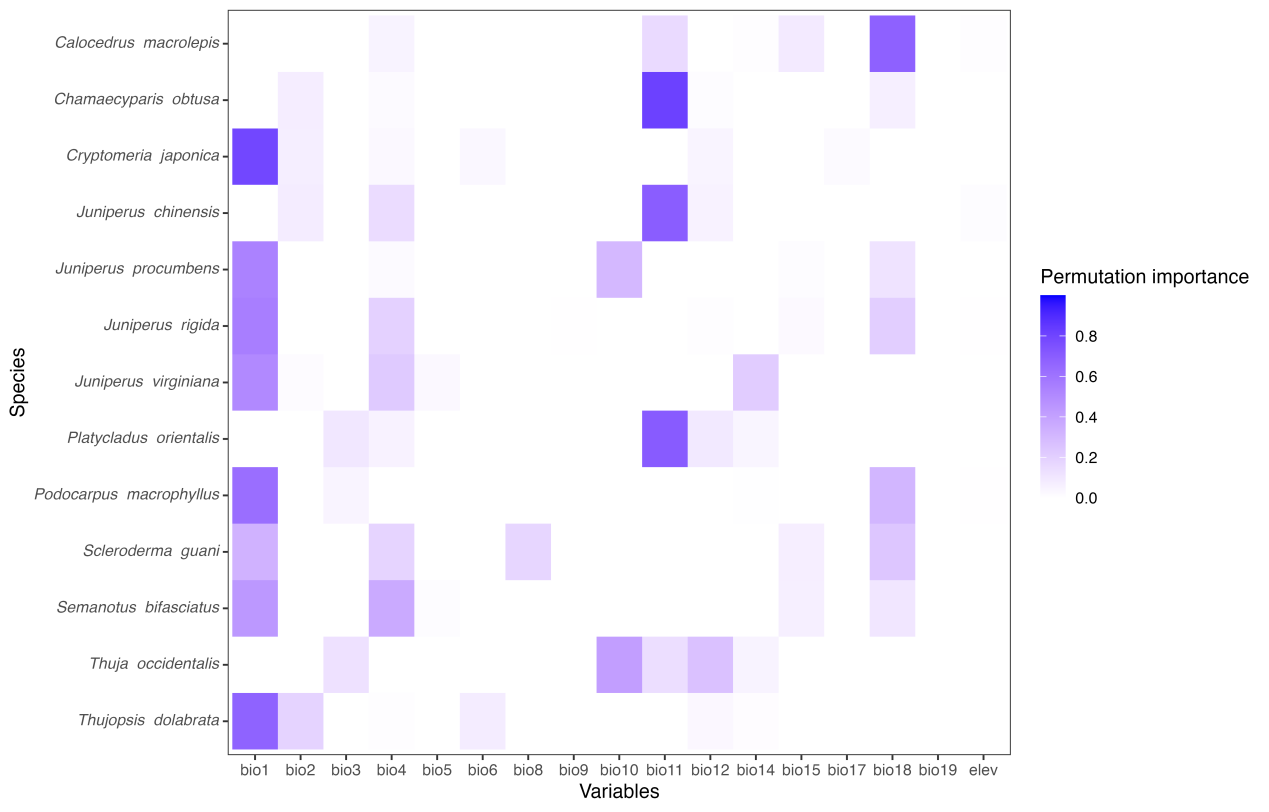

**Figure S2 :** Bioclimatic variable used in the projections for each species

The red gradient represents the variable performance importance value.

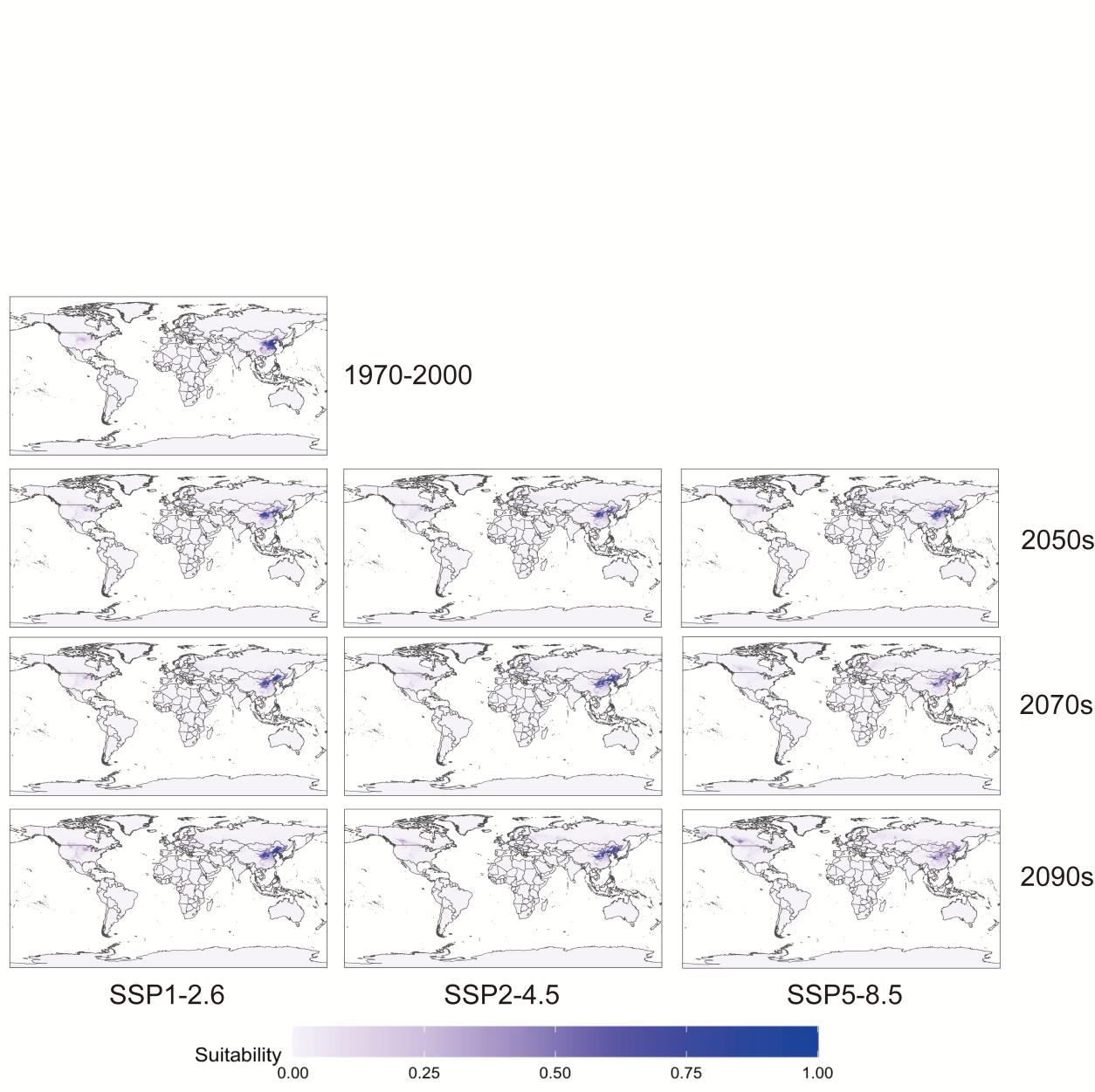

**Figure S3 :** The potential distribution of *Semanotus bifasciatus* under historical and future climates Blue gradient indicates the suitability of *Semanotus bifasciatus.*

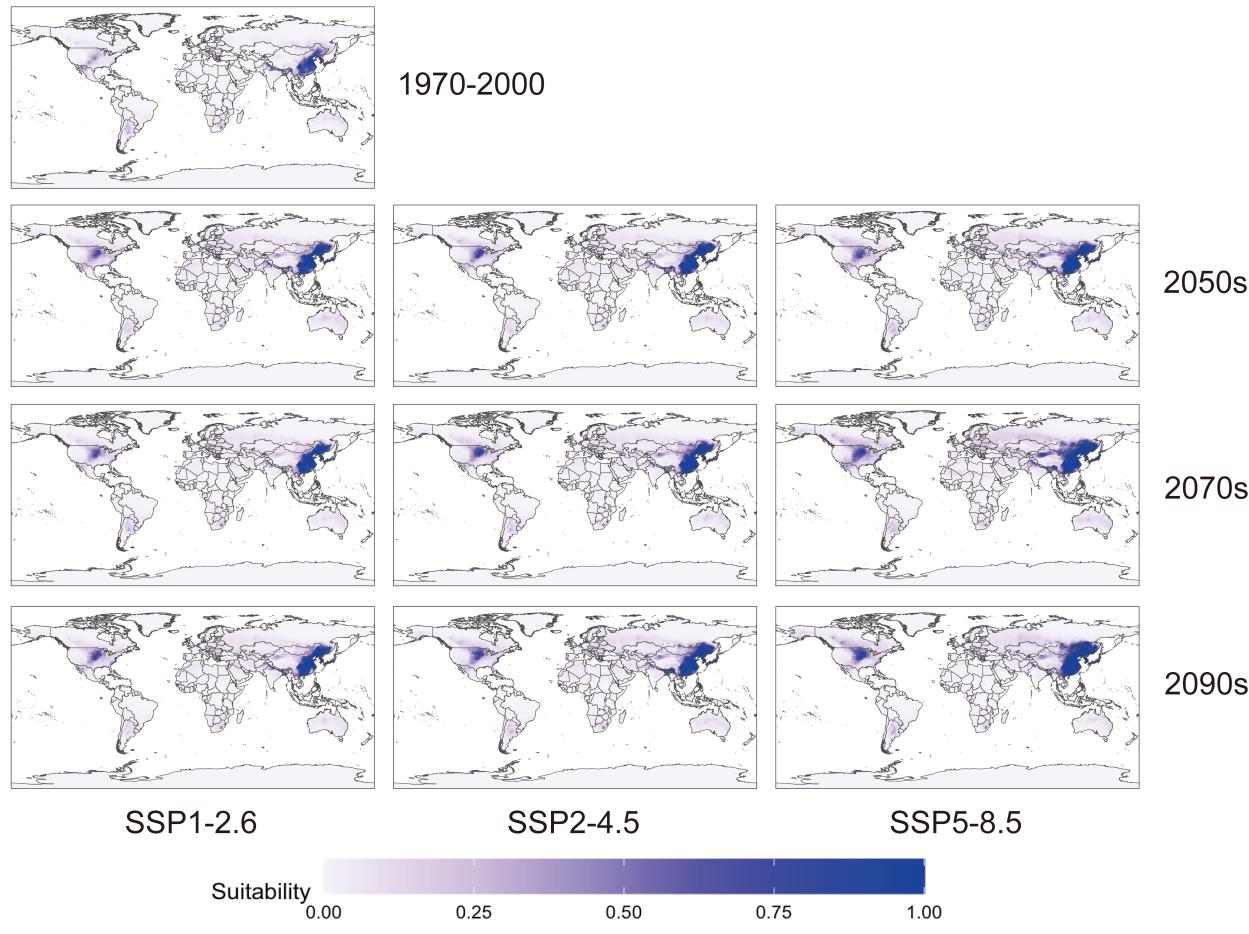

**Figure S4 :** The potential distribution of *Sclerodermus guani* under historical and future climates

Blue gradient indicates the suitability of *Sclerodermus guani.*

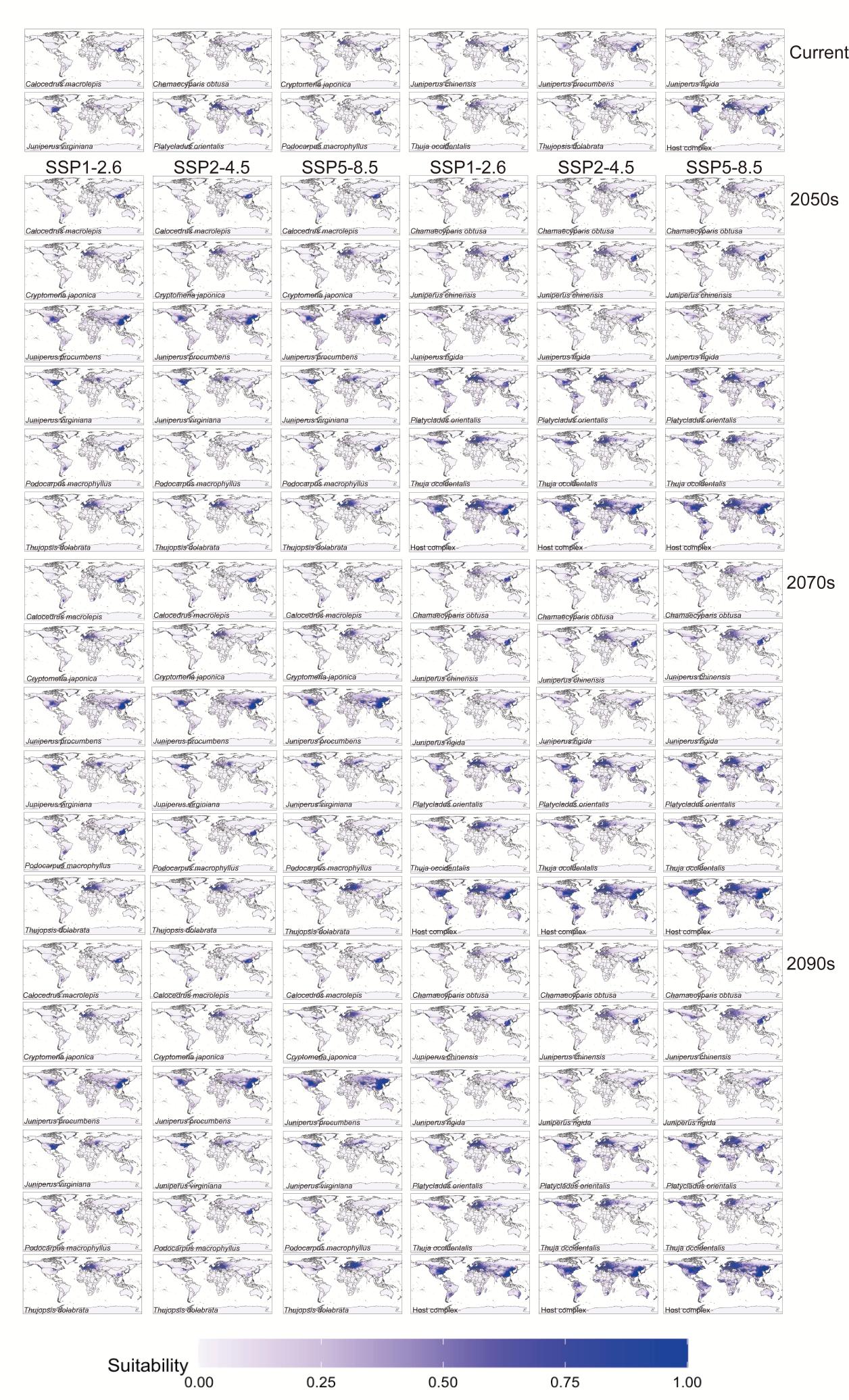

**Figure S5 :** The potential distribution of *Semanotus bifasciatus*’ 11 host plants and the host complex under historical and future climates

Blue gradient indicates the suitability of the host plants and the host complex.

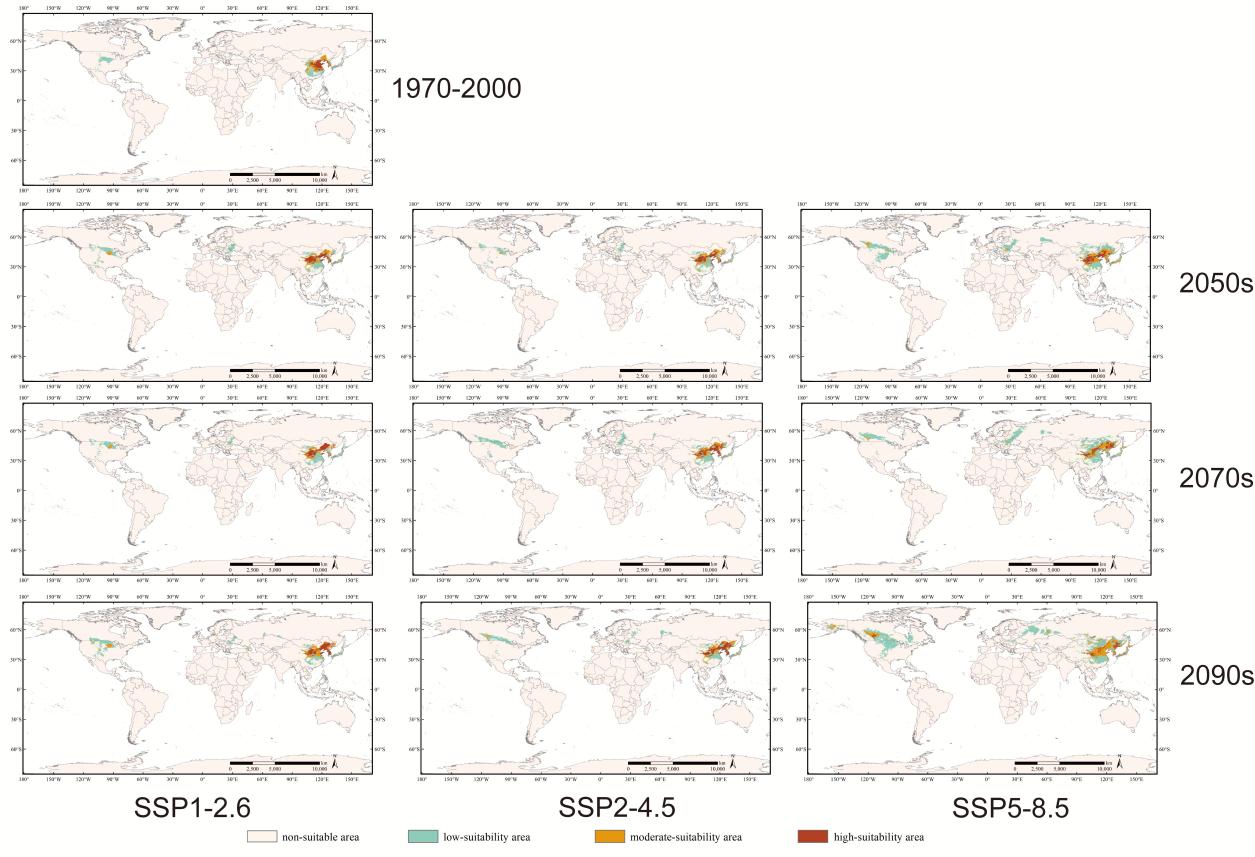

**Figure S6 :** The potential suitable area of *Semanotus bifasciatus* under historical and future climates predicted by the climate-only model

Beige indicates “non-suitable area”, green indicates “low-suitability area”, orange indicates “moderate-suitability area”, and red indicates “high-suitability area”.

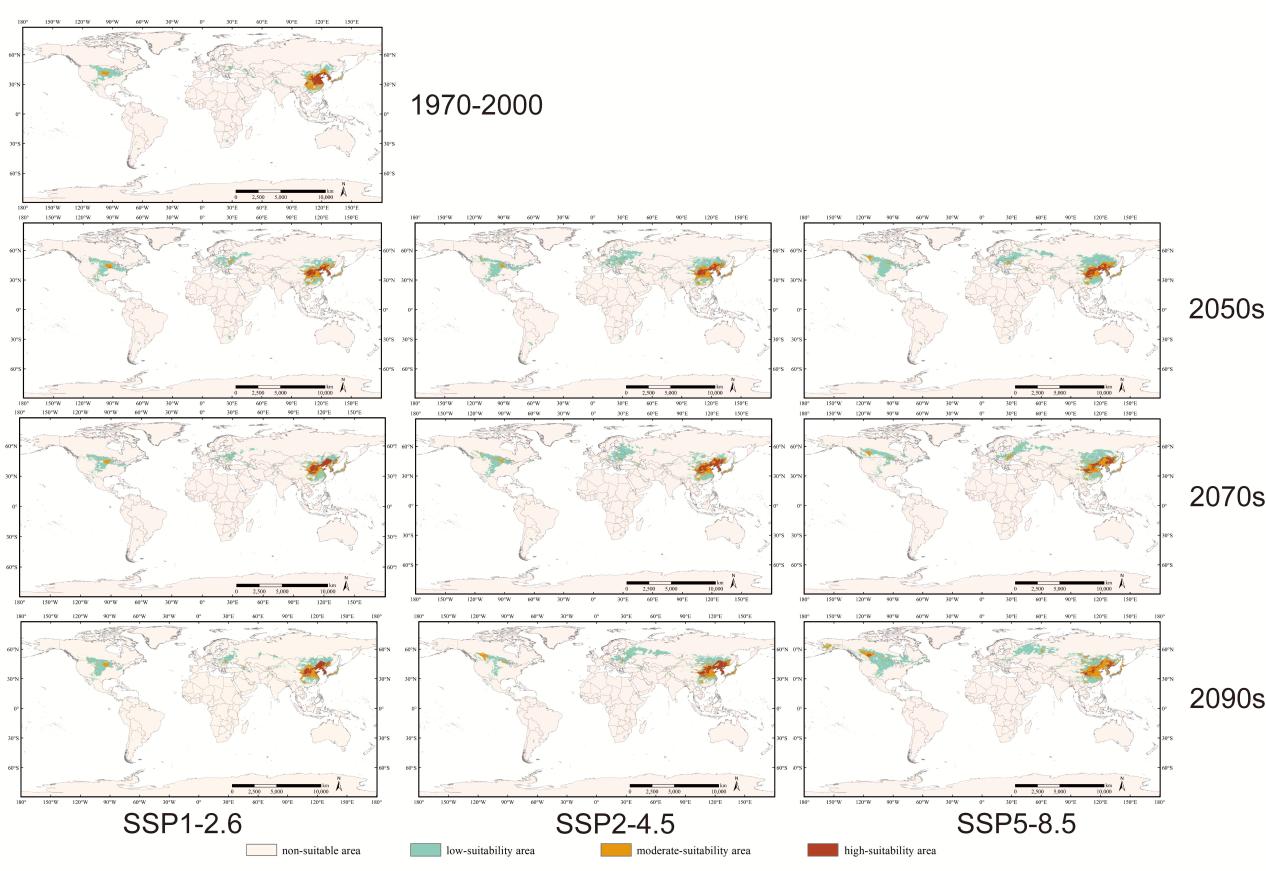

**Figure S7 :** The potential suitable area of *Semanotus bifasciatus* under historical and future climates predicted by the harmonic mean model

Beige indicates “non-suitable area”, green indicates “low-suitability area”, orange indicates “moderate-suitability area”, and red indicates “high-suitability area”.

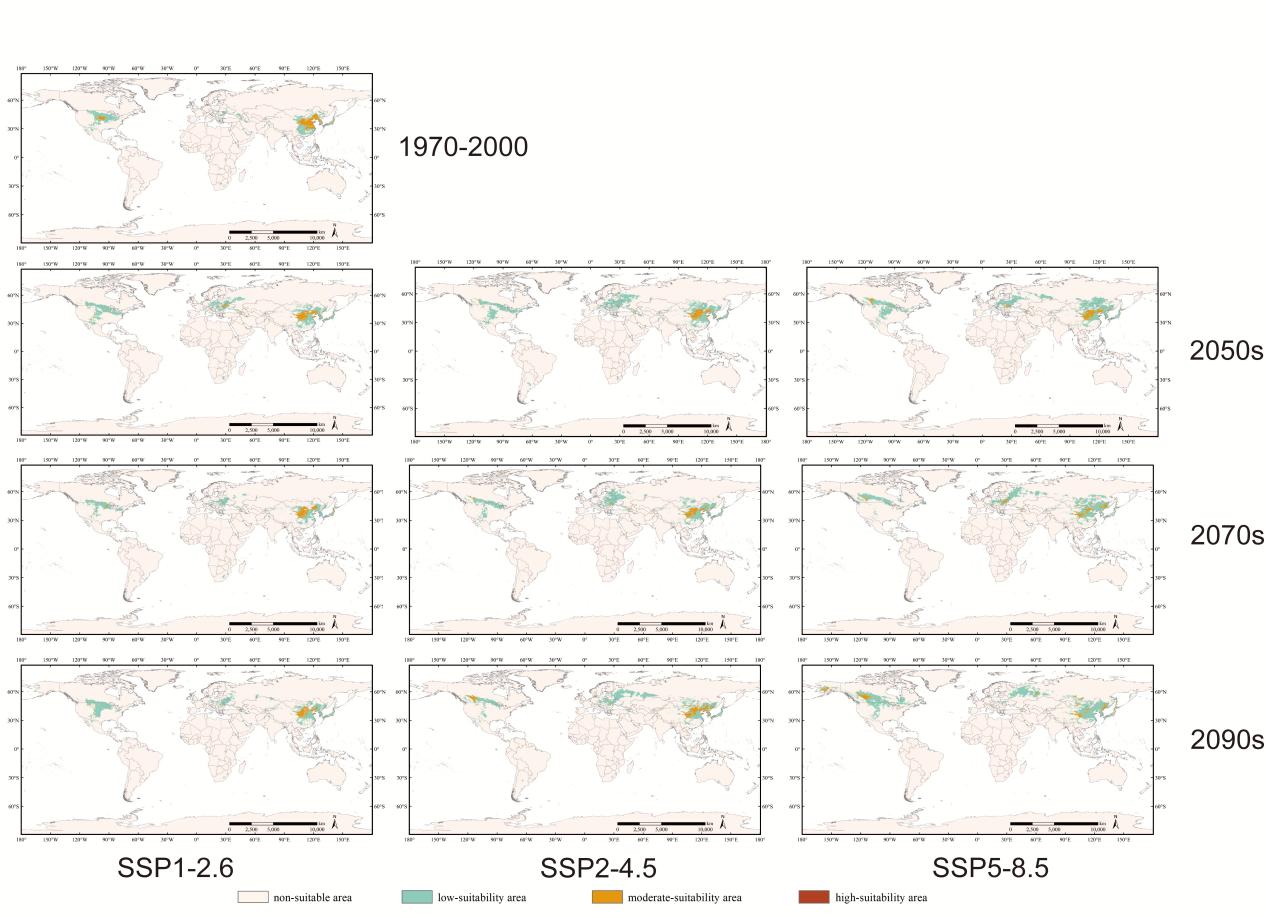

**Figure S8 :** The potential suitable area of *Semanotus bifasciatus* under historical and future climates predicted by the suppression model

Beige indicates “non-suitable area”, green indicates “low-suitability area”, orange indicates “moderate-suitability area”, and red indicates “high-suitability area”.

**Table S4 :** Areas of suitability classes (low, moderate, high) for each model, scenario, and period

| Model | Scenario | Period | Total suitable (10^6^ km²) | Low | Moderate | High |
| --- | --- | --- | --- | --- | --- | --- |
| Climate-only MaxEnt model | Current | 1970–2000 | 2.73 | 1.51 | 0.68 | 0.55 |
|  | SSP1-2.6 | 2050s | 4.05 | 2.41 | 0.93 | 0.72 |
|  |  | 2070s | 3.59 | 2.13 | 0.72 | 0.74 |
|  |  | 2090s | 4.24 | 2.37 | 0.99 | 0.88 |
|  | SSP2-4.5 | 2050s | 3.66 | 2.1 | 0.89 | 0.67 |
|  |  | 2070s | 4.27 | 2.73 | 0.78 | 0.77 |
|  |  | 2090s | 4.17 | 2.41 | 0.85 | 0.92 |
|  | SSP5-8.5 | 2050s | 6.57 | 4.53 | 1.27 | 0.77 |
|  |  | 2070s | 6.46 | 4.71 | 1.15 | 0.60 |
|  |  | 2090s | 9.80 | 6.87 | 2.40 | 0.53 |
| Harmonic mean model | Current | 1970–2000 | 5.49 | 3.26 | 1.31 | 0.89 |
|  | SSP1-2.6 | 2050s | 7.97 | 5.54 | 1.61 | 0.82 |
|  |  | 2070s | 6.56 | 4.53 | 1.14 | 0.89 |
|  |  | 2090s | 7.32 | 4.97 | 1.36 | 0.99 |
|  | SSP2-4.5 | 2050s | 9.77 | 7.32 | 1.64 | 0.81 |
|  |  | 2070s | 8.31 | 5.90 | 1.36 | 1.05 |
|  |  | 2090s | 8.27 | 5.55 | 1.52 | 1.19 |
|  | SSP5-8.5 | 2050s | 10.77 | 8.27 | 1.54 | 0.96 |
|  |  | 2070s | 10.05 | 7.36 | 1.88 | 0.81 |
|  |  | 2090s | 12.46 | 9.07 | 2.72 | 0.67 |
| Suppression model | Current | 1970–2000 | 4.82 | 3.42 | 1.40 | 0 |
|  | SSP1-2.6 | 2050s | 6.53 | 5.52 | 1.01 | 0 |
|  |  | 2070s | 4.99 | 4.09 | 0.90 | 0 |
|  |  | 2090s | 5.71 | 4.88 | 0.83 | 0 |
|  | SSP2-4.5 | 2050s | 8.07 | 7.11 | 0.96 | 0 |
|  |  | 2070s | 6.67 | 5.80 | 0.87 | 0 |
|  |  | 2090s | 6.58 | 5.34 | 1.25 | 0 |
|  | SSP5-8.5 | 2050s | 8.79 | 7.72 | 1.06 | 0 |
|  |  | 2070s | 7.73 | 6.83 | 0.88 | 0.02 |
|  |  | 2090s | 9.79 | 8.65 | 1.3 | 0.04 |

**Table S5 :** Centroid locations and migration distance of suitable areas for *Semanotus bifasciatus* under three modes

| Model | Region | Scenario | Current centroid (lat, lon) and Direction | 2050s centroid | Direction | Distance (km) |
| --- | --- | --- | --- | --- | --- | --- |
| Climate-only MaxEnt model | Asia | SSP1-2.6 | 35.802°N,114.215°E  Hebi, Henan, China | 36.897°N,106.182°E | Wuzhong, Ningxia, China | 729.42 |
|  |  | SSP2-4.5 |  | 37.970°N,107.851°E | Ordos, Inner Mongolia Autonomous Region, China | 615.02 |
|  |  | SSP5-8.5 |  | 40.212°N,104.914°E | Alxa Right Banner, Alxa League, Inner Mongolia Autonomous Region, China | 950.29 |
|  | North America | SSP1-2.6 | 36.978°N,103.865°W  Union, New Mexico, United States | 43.238°N,107.571°W | Fremont, Wyoming, United States | 763.90 |
|  |  | SSP2-4.5 |  | 48.048°N,113.172°W | Tooele, Utah, United States | 1445.73 |
|  |  | SSP5-8.5 |  | 53.365°N,120.880°W | Fraser-Fort George, British Columbia, Canada | 2245.78 |
|  | Europe | SSP1-2.6 | 45.272°N,35.095°E  Crimea, Ukraine | 45.936°N,28.591°E | Balabanu, Taraclia, Moldova | 511.18 |
|  |  | SSP2-4.5 |  | 53.151°N,34.511°E | Bryansky District, Bryansk Oblast, Russia | 877.12 |
|  |  | SSP5-8.5 |  | 54.151°N,35.778°E | Kozel'skiy rayon, Kaluga Oblast, Russia | 988.51 |
| Harmonic mean model | Asia | SSP1-2.6 | 34.207°N,109.813°E  Shangluo, Shaanxi, China | 37.494°N,103.560°E | Wuwei, Gansu, China | 671.47 |
|  |  | SSP2-4.5 |  | 38.367°N,105.479°E | Alxa Left Banner, Alxa League, Inner Mongolia Autonomous Region, China | 603.88 |
|  |  | SSP5-8.5 |  | 40.956°N,104.509°E | Alxa Left Banner, Alxa League, Inner Mongolia Autonomous Region, China | 883.67 |
|  | North America | SSP1-2.6 | 37.186°N,99.646°W  Clark, Kansas, United States | 43.552°N,107.379°W | Washakie, Wyoming, United States | 963.65 |
|  |  | SSP2-4.5 |  | 47.310°N,110.568°W | Judith Basin, Montana, United States | 1437.74 |
|  |  | SSP5-8.5 |  | 53.371°N,119.408°W | Division No. 15, Alberta, Canada | 2356.49 |
|  | Europe | SSP1-2.6 | 45.964°N,24.105°E  Sibiu County, Romania | 48.329°N,26.845°E | Coteala, Briceni, Moldova | 334.76 |
|  |  | SSP2-4.5 |  | 50.246°N,28.709°E | Kosmonavtov street, Zhytomyr Oblast, Ukraine | 585.90 |
|  |  | SSP5-8.5 |  | 54.291°N,33.792°E | Spas-Demensky District, Kaluga Oblast, Russia | 1152.84 |
| Suppression model | Asia | SSP1-2.6 | 35.40°N,108.46°E  Qingyang, Gansu, China | 38.69°N,100.31°E | Zhangye, Gansu, China | 810.14 |
|  |  | SSP2-4.5 |  | 39.79°N,101.19°E | Alxa Right Banner, Alxa League, Inner Mongolia Autonomous Region, China | 804.85 |
|  |  | SSP5-8.5 |  | 41.37°N,100.98°E | Ejina Banner, Alxa League, Inner Mongolia Autonomous Region, China | 929.72 |
|  | North America | SSP1-2.6 | 37.45°N,99.81°W  Clark, Kansas, United States | 43.85°N,109.51°W | Fremont, Wyoming, United States | 1083.14 |
|  |  | SSP2-4.5 |  | 47.75°N,111.88°W | Teton, Montana, United States | 1508.90 |
|  |  | SSP5-8.5 |  | 53.14°N,118.49°W | Division No. 15, Alberta, Canada | 2261.59 |
|  | Europe | SSP1-2.6 | 45.94°N,24.10°E  Sibiu County, Romania | 47.98°N,26.66°E | Satu Mare County, Romania | 298.63 |
|  |  | SSP2-4.5 |  | 49.83°N,28.77°E | Vinnytsia Oblast, Ukraine | 555.08 |
|  |  | SSP5-8.5 |  | 54.58°N,34.59°E | Mosal'sky District, Kaluga Oblast, Russia | 1213.47 |
